# Endoplasmic reticulum (ER) lumenal indicators in *Drosophila* reveal effects of HSP-related mutations on ER calcium dynamics

**DOI:** 10.1101/2020.02.20.957696

**Authors:** Megan K Oliva, Juan José Perez-Moreno, Jillian O’Shaughnessy, Trevor J Wardill, Cahir J O’Kane

## Abstract

Genes for ER-shaping proteins are among the most commonly mutated in Hereditary Spastic Paraplegia (HSP). Mutation of these genes in model organisms can lead to disruption of the ER network. To investigate how the physiological roles of the ER might be affected by such disruption, we developed tools to interrogate its Ca^2+^ signaling function. We generated GAL4-driven Ca^2+^ sensors targeted to the ER lumen, to record ER Ca^2+^ fluxes in identified *Drosophila* neurons. Using *GAL4* lines specific for Type Ib or Type Is larval motor neurons, we compared the responses of different lumenal indicators to electrical stimulation, in axons and presynaptic terminals. The most effective sensor, ER-GCaMP6-210, had a Ca^2+^ affinity close to the expected ER lumenal concentration. Repetitive nerve stimulation generally showed a transient increase of lumenal Ca^2+^ in both the axon and presynaptic terminals. Mutants lacking neuronal reticulon and REEP proteins, homologs of human HSP proteins, showed a larger ER lumenal evoked response compared to wild type; we propose mechanisms by which this phenotype could lead to neuronal dysfunction or degeneration. Our lines are useful additions to a *Drosophila* Ca^2+^ imaging toolkit, to explore the physiological roles of ER, and its pathophysiological roles in HSP and in axon degeneration more broadly.

## Introduction

Hereditary Spastic Paraplegia (HSP) is a genetically heterogeneous disorder, with over 80 loci and 60 genes identified. It also shows phenotypic heterogeneity in manifestations such as age of onset, and the presence of other symptoms (Blackstone 2018). Despite this heterogeneity, there is evidence for some common mechanisms of cellular pathophysiology. The endoplasmic reticulum (ER) is implicated in these mechanisms, with HSP mutations affecting ER proteins with a variety of roles including lipid metabolism, ER membrane-contact-site function, and ER architecture (Blackstone 2018). Gene products that shape the tubular ER network are among the most commonly mutated in HSP, and mutation of these genes in model organisms leads to physical disruption of the ER network (O’Sullivan et al. 2012, Fowler and O’Sullivan 2016, Summerville et al. 2016, Yalçın et al. 2017, Lindhout et al. 2019). These studies point to the importance of a continuous and connected tubular ER network throughout axons and presynaptic terminals, and suggest disruption to this network as a common thread in HSP pathogenesis. Although changes in synaptic structure and function are found in these mutants (Summerville et al. 2016, Li et al. 2017, Lindhout et al. 2019), the direct physiological consequences of altered ER architecture remain elusive. It is imperative to understand the roles of ER architecture and the consequences of disrupting it, to understand how this might result in a neurodegenerative cascade.

We therefore need tools to study the physiological functions of axonal and presynaptic ER that depend on its architecture. Recently, de Juan-Sanz et al (2017) developed sensitive Genetically Encoded Calcium Indicators (GECIs) targeted to the ER lumen, to examine the role of the ER in Ca^2+^ handling and signaling in synaptic activity in mammalian cultured neurons. Here we have expressed one of these probes (ER-GCaMP6-210) under GAL4 control in *Drosophila* motor neurons, and targeted two other Ca^2+^ sensors with higher Ca^2+^ affinities, CEPIA3-ER and CEPIA4-ER (Suzuki et al. 2014) to the *Drosophila* ER lumen. We show ER-GCaMP6-210 as the most effective reporter of the three by consistently producing the largest changes in fluorescent intensity, and characterize its localization and its reporting of ER behavior in axon and presynaptic terminals. We then use it to show perturbation of Ca^2+^ handling in *Drosophila* in which ER organization is disrupted by loss of HSP-related ER-modeling proteins.

## Materials and Methods

### Gene Constructs

The *pUASTattB* vector (Bischof et al. 2007), was previously modified to include *attR* sites and the *ccdB* gene to make it compatible with the Gateway cloning system (Invitrogen) (Moreau et al. 2014). We further modified this vector, increasing the number of GAL4-binding sites from 5 to 17, to generate *p17xUASTattB*, and increase GAL4-dependent gene expression (Supplementary File 1). To generate *p17xUASTattB-ER-GCaMP6-210*, we PCR-amplified a fragment encoding GCaMP210-KDEL fused to a calreticulin signal peptide, from Addgene plasmid *ER-GCaMP6-210* (de Juan-Sanz et al. 2017) (Table 1), while adding *attB1* and *attB2* sites to its 5’ and 3’ ends to enable it to integrate into *p17xUASTattB* using the Gateway cloning system (Supplementary File 1). To generate *p17xUASTattB-CEPIA3-ER* and *p17xUASTattB-CEPIA4-ER*, we PCR-amplified the CEPIA coding region from Addgene plasmids *pCMV CEPIA3mt* and *pCMV CEPIA4mt* (Suzuki et al. 2014) (Table 1), while adding *attB1* and a BiP signal sequence to the 5’ end, and HDEL and *attB2* to the 3’ end (BiP and HDEL ER-targeting as in (Summerville et al. 2016)), and then integrating it into *p17xUASTattB* (Supplementary File 1).

**Table 1:** Stocks and plasmids used in this work.

| **Drosophila Stock Genotype** | **RRID** | **Source/Reference** |
| --- | --- | --- |
| *y w; +; 17xUASTattB-ER-GCaMP6-210(attP86Fb)* | - | This paper |
| *y w; +; 17xUASTattB-CEPIA3-ER(attP86Fb)* | - | This paper |
| *y w; +; 17xUASTattB-CEPIA4-ER(attP86Fb)* | - | This paper |
| *w<sup>1118</sup>; +; UAS-tdTomato::Sec61β(attP2)* | BDSC_64747 | (Summerville et al. 2016) |
| *w; UAS-Sturkopf::GFP; +* | - | (Thiel et al. 2013) |
| *w; +; UAS-jRCaMP1b(VK00005)* | BDSC_63793 | (Dana et al. 2016) |
| *w; Rtnl1<sup>-</sup> ReepA<sup>-</sup> ReepB<sup>-</sup>/CyO; +* | BDSC_77904 | (Yalçın et al. 2017) |
| *w; ReepA<sup>+</sup>; + (WT control)* | - | (Yalçın et al. 2017) |
| *w<sup>1118</sup>; +; Dpr-GMR94G06-GAL4(attP2)* | BDSC_40701 | (Pfeiffer et al. 2008, Perez-Moreno and O'Kane 2019) |
| *w<sup>1118</sup>; +; FMR-GMR27E09-GAL4(attP2)* | BDSC_49227 | (Pfeiffer et al. 2008, Perez-Moreno and O'Kane 2019) |
| **Plasmid** |  |  |
| *ER-GCaMP6-210* | Addgene_86919 | Tim Ryan (de Juan-Sanz et al. 2017) |
| *pCMV CEPIA3mt* | Addgene_58219 | Masamitsu Iino (Suzuki et al. 2014) |
| *pCMV CEPIA4mt* | Addgene_58220 | Masamitsu Iino (Suzuki et al. 2014) |
| *p17xUASTattB* | - | This paper |
| *p17xUASTattB-ER-GCaMP6-210* | - | This paper |
| *p17xUASTattB-CEPIA3-ER* | - | This paper |
| *p17xUASTattB-CEPIA4-ER* | - | This paper |

### *Drosophila* Genetics

*p17xUASTattB-ER-GCaMP6-210, p17xUASTattB-CEPIA3-ER* and *p17xUASTattB-CEPIA4-ER* were injected into *Drosophila* embryos and integrated at landing site *attP86Fb* (Bischof et al. 2007) using phiC31 integrase-mediated integration, by the University of Cambridge Genetics Department injection service. Other fly stocks used can be found in Table 1.

### Histology

Third instar larvae were dissected in ice-cold Ca^2+^-free HL3 solution (Stewart et al. 1994). HL3 was then replaced with PBS and larvae fixed for 10 min in PBS with 4% formaldehyde. Fixed preparations were mounted in Vectashield (Vector Laboratories), and images were collected using EZ-C1 acquisition software (Nikon) on a Nikon Eclipse C1si confocal microscope (Nikon Instruments, UK). Images were captured using a 40x/1.3NA oil objective.

### Live Imaging

Third instar larvae were dissected in ice-cold Schneider’s insect medium (Sigma). Before imaging, Schneider’s medium was replaced with HL3 containing (in mM) 70 NaCl, 5 KCl, 0.5 MgCl_2_, 10 NaHCO_3_, 115 sucrose, 5 trehalose, 5 HEPES, 1 CaCl_2_ and 7 L-glutamic acid, pH ∼7.2 ± 0.05. The ER lumenal response to electrical stimulation was sensitive to Mg^2+^, with only very small or no responses elicited in HL3 containing 20 mM MgCl_2_, which was used for fixed samples; a low-Mg^2+^ HL3 solution (0.5 mM MgCl_2_) gave much more reliable responses.

Segmental nerves were cut with dissection scissors and then drawn by suction into a heat-polished glass pipette, ∼10 μm internal diameter (Macleod et al. 2002), which was connected to a stimulator and train generator (DS2A-MK.II Constant Voltage Stimulator and DG2A Train/Delay Generator (Digitimer, Welwyn Garden City, UK)) to deliver supra-threshold electrical pulses (∼5 V). Larval NMJ responses were recorded in abdominal segments A4-A6. After 10 s of baseline image acquisition, 400-μs impulses were delivered for 2 s at a range of frequencies. The interval between different stimulation frequencies was ∼60 seconds, or until ER lumen fluorescence returned to baseline levels (whichever was longer).

Wide-field Ca^2+^ imaging was performed on an upright Zeiss Axiskop2 microscope with Optosplit II (Cairn) using a 40x NA 1.0 water-immersion objective (W Plan-Apochromat 40x/1.0 DIC M27), a 2x C-Mount Fixed Focal Length Lens Extender (Cairn, Faversham, UK) and an Andor EMCCD camera (Model iXon Ultra 897_BV, 512×512 pixels, Andor Technology, Belfast, UK) at 10 frames per second, 100 ms exposure, and EM gain level 100. Imaging data were acquired using Micro-Manager (Edelstein et al. 2014) and saved as multi-layer tif files before analysis using ImageJ FIJI (https://fiji.sc) (Schindelin et al. 2012).

### Image analysis and Figure Preparation

Confocal microscopy images of fixed samples were collected using EZ-C1 acquisition software on a Nikon Eclipse C1si confocal microscope. Images were analyzed and processed using ImageJ Fiji (https://fiji.sc) (Schindelin et al. 2012). Wide-field live images were opened in ImageJ Fiji, and channels were individually bleach-corrected (simple ratio, background intensity=0). Fluorescent intensity time courses were obtained for an ROI traced around the entire NMJ, or axon segment, in each frame in a given data set using Time Series Analyzer V3 plugin, saved as txt files and fed into R scripts written by the authors to obtain the ΔF/F data (Supplementary File 2). Graphs were generated using GraphPad Prism 8. Figures were made using Adobe Illustrator.

### Statistical analysis

Data were analyzed in GraphPad Prism, using 2-way ANOVA, unpaired two-tailed Student’s t-tests; graphs show mean ± SEM, or Mann-Whitney U tests (for data not normally distributed); graphs show median with interquartile range. Pearson’s correlation was used to examine relationships between datasets. Sample sizes are reported in figures.

## Results

### The ER lumenal Ca^2+^ evoked response is more delayed and sustained compared to the cytoplasmic response

To monitor Ca^2+^ flux in the ER lumen, we generated transgenic flies that encoded ER-GCaMP6-210 (de Juan-Sanz et al. 2017), a GCaMP6 sensor fused to a signal peptide and ER retention signal, with a Ca^2+^ affinity optimized for the relatively high [Ca^2+^] in ER lumen. ER-GCaMP6-210 localization in presynaptic terminals largely overlapped with the ER marker tdTomato::Sec61β (Fig. 1A), being more restricted than cytoplasmic jRCaMP1b (Fig. 1B), consistent with localization to the ER lumen. To test the response of the sensor to neuronal activation we electrically stimulated the nerve. The cytoplasmic Ca^2+^ response quickly reached a peak in the first couple of seconds following stimulation, and decayed back to resting levels over several seconds. In contrast, the ER-GCaMP6-210 lumenal response increased more slowly, reaching a peak five or more seconds following stimulation, with a slow return to resting levels, in some cases over minutes (Fig. 2A,B; Supplementary Video 1), in keeping with the slower dynamics reported previously for these constructs (de Juan-Sanz et al. 2017). Cytoplasmically jRCaMP1b and GCaMP6f have similar kinetics in neurons as reported previously, with just a slightly slower half decay time (0.6s) for jRCaMP1b (Dana et al. 2016), compared to GCaMP6f (0.4s) (Chen et al. 2013) at 10 Hz. In around 10% of nerves, usually at higher stimulation frequencies, we alternatively (or sometimes additionally) observed small, sharp decreases in ER lumenal fluorescence immediately following stimulation. This decrease was followed by a quick return to baseline fluorescence, or by a slow increase in fluorescence as described above. This short-term decrease in fluorescence was observed more frequently in the axon than at the NMJ. There did not appear to be anything notably different in the cytoplasmic flux in these instances, compared to when the ER flux showed only an increase in fluorescence (Fig. 2A; Fig. S1). Together this data suggests a slower and somewhat less predictable ER Ca^2+^ flux, compared to the cytoplasmic flux in axons and NMJs of motor neurons.

**Fig 1.**
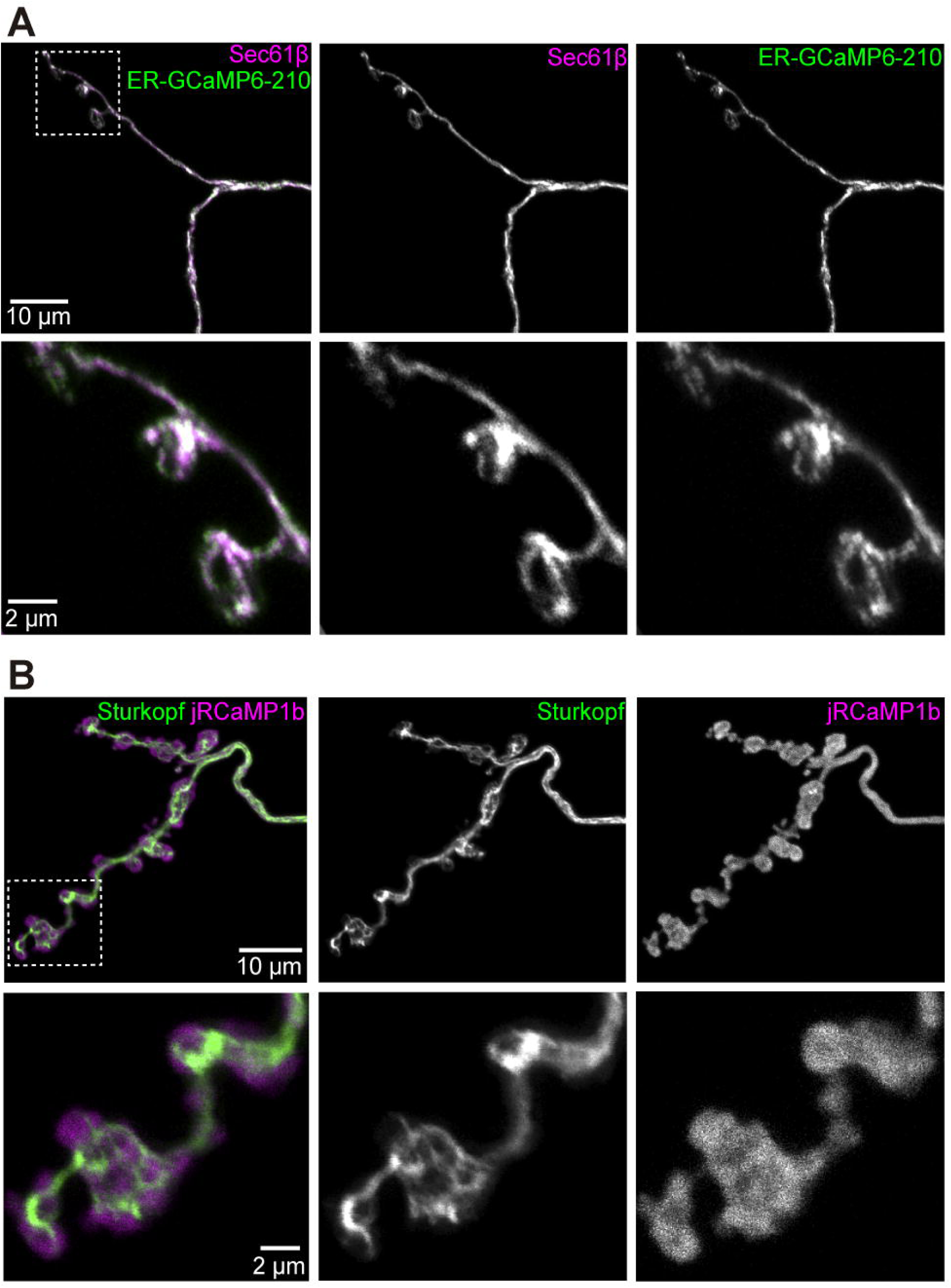
The ER lumenal sensor ER-GCaMP6-210 has an overlapping distribution with ER at the neuromuscular junction. Panels show markers in Ib boutons at muscle 1 NMJ, segment A2, expressed using *Dpr-GMR94G06-GAL4*; all images are maximum intensity projections of confocal stacks. (**A**) Overlap of ER-GCaMP6-210 and ER membrane marker tdTomato::Sec61β. (**B**) jRCaMP1b is more broadly distributed than ER membrane marker Sturkopf::GFP. Magnified views of each inset are in respective lower rows.

**Fig 2.**
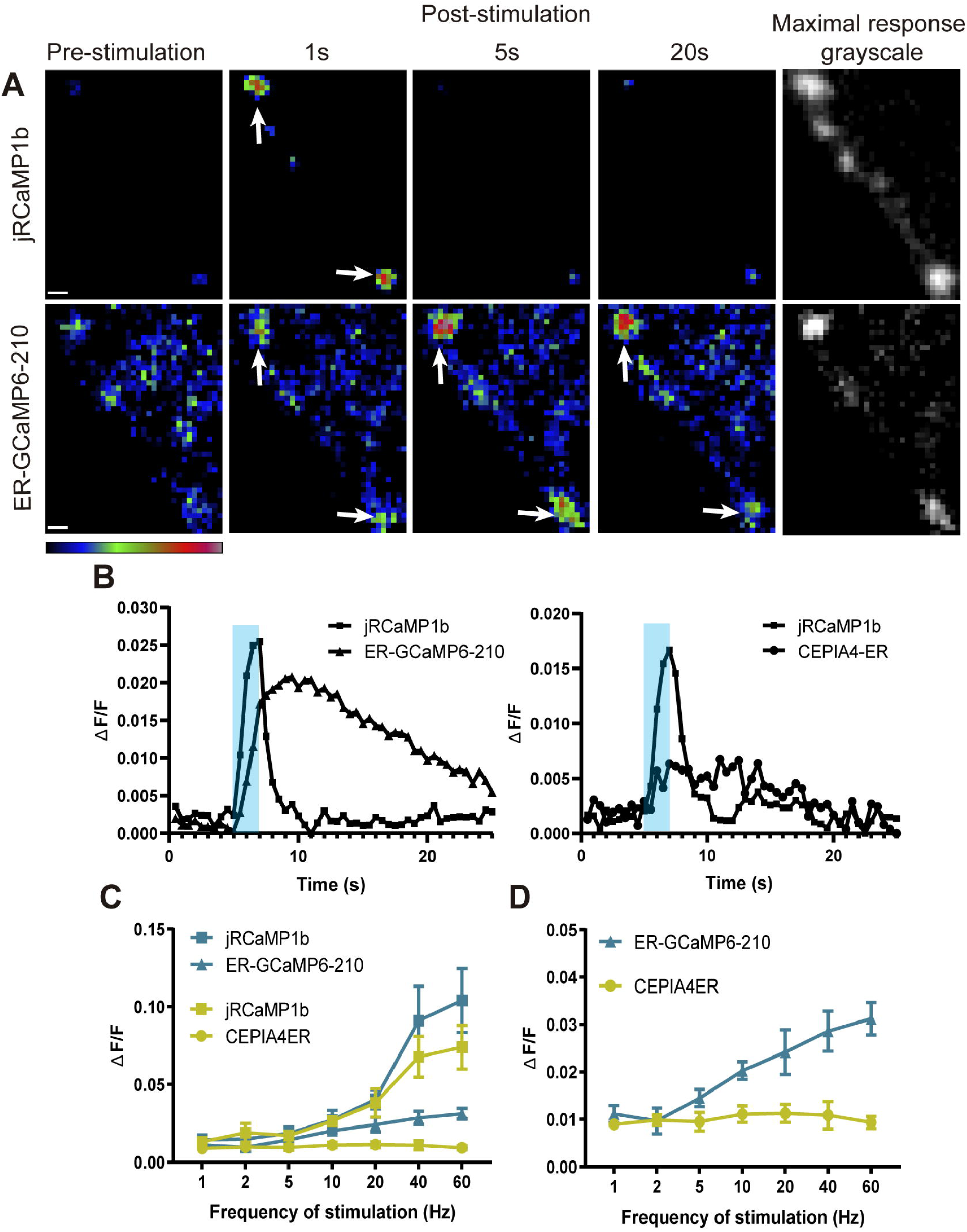
ER lumenal Ca^2+^ responses to electrical stimulation at the NMJ, visualized using ER-GCaMP6-210, were delayed and more sustained compared to cytoplasmic responses. (**A**) Representative pseudocolour images of responses to 40 Hz stimulation, with maximal responses shown in grayscale (right panels) to show position of synapse. Panels show responses of jRCaMP1b (top row) and ER-GCaMP6-210 (bottom row) at muscle 6/7, expressed in the same RP5 Type Is motor neuron using *FMR-GMR27E09-GAL4*. Arrows show examples of boutons. Pseudocolour bar shows low to high relative intensity; scale bar, 2 µm. (**B**) Representative responses to 20 Hz stimulation, of ER-GCaMP6-210 and CEPIA4-ER with corresponding jRCaMP1b response at muscle 6/7, expressed in RP5 Type Is motor neuron using *FMR-GMR27E09-GAL4*. Shading represents the stimulation period. (**C**) Average fold-change in peak fluorescence at different stimulation frequencies, of ER-GCaMP6-210 (green) and CEPIA4-ER, compared with jRCaMP1b (magenta) at muscle 6/7, expressed in the same RP5 Type Is motor neuron using *FMR-GMR27E09-GAL4*. (ER-GCaMP6-210, N=5; CEPIA4-ER, N=6). (**D**) Comparison of ER lumenal sensors only, replotted from (**C**) with the y-axis expanded.

### ER lumenal sensor with affinity closest to predicted ER lumenal [Ca^2+^] shows the strongest response

Since the high surface/volume ratio of the small ER tubules of axons might potentially make them more susceptible than other ER regions to leakage of Ca^2+^ to the cytosol, we generated transgenic flies carrying two additional ER lumenal sensors with higher Ca^2+^ affinities. The highest affinity sensor, CEPIA3-ER (K_d_=11 µM), showed very small or no responses at multiple NMJs of seven larval preparations, even when a robust cytoplasmic response was elicited (not shown); CEPIA4-ER (K_d_=59 µM) usually showed a small response at most frequencies. ER-GCaMP6-210 (K_d_=210 µM) produced the most consistent response, and the magnitude of its response increased with increasing stimulation frequency (Fig. 2B-D). As sensors with a K_d_ that approximates the surrounding [Ca^2+^] are best suited as reporters, this is consistent with the estimated resting neuronal ER lumenal [Ca^2+^] of 150-200 µM reported previously (de Juan-Sanz et al. 2017).

### The axonal cytoplasmic evoked response is smaller compared to NMJ response, whereas the ER lumenal evoked response is consistent across locations

As we were interested to see if we could observe Ca^2+^ signals along the length of the axon, we used *FMR-GMR27E09-GAL4* to record in the two Type Is motor neurons per hemisegment where it drives expression: RP2 and RP5, which together innervate several NMJs (Perez-Moreno and O’Kane 2019). We compared several axonal and NMJ locations within a muscle hemisegment (Fig. S2A). Motor neuron RP2 (also known as ISN-Is) innervates dorsal muscles, and we recorded from its NMJ boutons at muscle 1, and from two axonal locations: the “entry” point at which the axon starts to cross the muscles as it emerges from the ventral nerve cord, and more laterally as it passes over muscle 4. Motor neuron RP5 (also known as ISN b/d) innervates ventral muscles, and we recorded from its NMJ at muscle 6/7, and from one axonal location, the “entry” point, as defined for motor neuron RP2 (Fig. S2A,B; Supplementary Video 2). *Dpr-GMR94G06-GAL4* drives expression in the aCC Type Ib motor neuron (also known as MN1-Ib), which innervates muscle 1 (Perez-Moreno and O’Kane 2019). The responses of both cytosolic and lumenal sensors at the aCC muscle 1 NMJ boutons were comparable to those in RP2 at the same muscle (Fig. 3A,B), suggesting a comparable action potential response between the two neuronal types. Fluorescence of the sensors driven by *Dpr-GMR94G06-GAL4* in the axons of the Type Ib aCC neuron was too dim to obtain reliable evoked responses, which may be due to a relatively small diameter of its axon, and hence relatively low amounts of sensors at this subcellular location.

**Fig 3.**
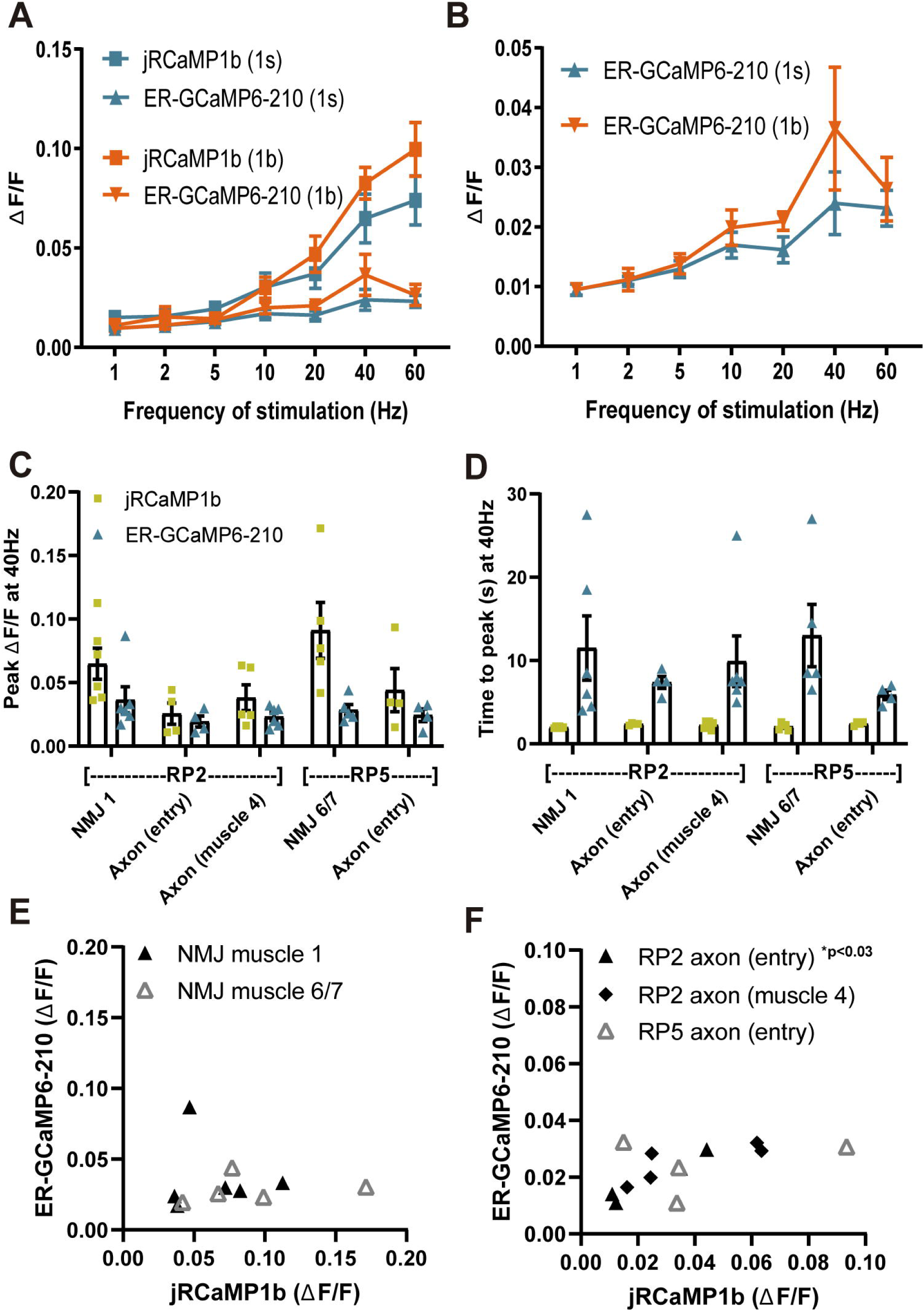
Comparisons of ER lumenal and cytoplasmic Ca^2+^ responses across axons and NMJs of Type Ib and Is motor neurons. (**A**) Evoked responses of lumenal ER-GCAMP6-210 and cytoplasmic jRCaMP1b to a range of stimulation frequencies, in Type Ib boutons of the aCC neuron at the muscle 1 NMJ, expressed using *Dpr-GMR94G06-GAL4* (N=6), or in Type Is boutons at the muscle 1 NMJ of the RP2 neuron expressed using *FMR-GMR27E09-GAL4* (N=6). (**B**) Comparison of ER lumen sensor data only, from (**A**) but with y-axis expanded. (**C**) Lumen and cytoplasmic Ca^2+^ fluxes at different positions of Type Is neurons RP2 and RP5 (Fig. S2A), with sensors expressed using *FMR-GMR27E09-GAL4*. (**D**) The time to peak ΔF/F, for both lumen and cytoplasmic responses, was similar across all axon and NMJ positions. (**E, F**) Peak amplitudes (from **C**) of individual ER lumen and cytoplasm responses were plotted for the NMJ (**E**) and axonal (**F**) compartments of RP2 and RP5. A positive correlation was found only for RP2 axon entry but not for other locations (Table 2). One outlier in RP2 axon (muscle 4) jRCaMP1b was excluded from graphs (**C**) and (**F**) (value 0.29), but included in analysis in (**F**).

The evoked cytoplasmic NMJ and axonal Ca^2+^ responses were both comparable between the two Type Is motor neurons, although axonal responses were consistently smaller than NMJ responses whereas ER lumenal evoked responses were consistent across all NMJ and axonal locations (Fig. 3C). This did not appear to be due to alterations in ER volume as resting fluorescence was consistent across locations (Fig. S2C). Both cytoplasmic and ER lumenal responses reached peak fluorescence on similar timescales across locations (Fig. 3D), suggesting that the observed responses were due to Ca^2+^ flux across the plasma membrane, and not a propagation of Ca^2+^ down the axon. The ER lumenal response appeared in general not to be influenced by the cytoplasmic response; most locations showed no correlation of peak ΔF/F between ER lumen and cytosol, apart from RP2 axon at entry point (Fig. 3E,F, Table 2). This suggests a more complex transfer of Ca^2+^ within and between intracellular compartments, than simply a flow of Ca^2+^ from cytosol to ER lumen.

**Table 2:** Correlation analysis of peak ΔF/F between ER lumen and cytosol in NMJ and axons using Pearson’s correlation coefficient.

| **FMR-GMR27E09 Neuronal location** | **r** | **p** | **N (larvae)** |
| --- | --- | --- | --- |
| RP2 NMJ muscle 1 | 0.10 | 0.85 | 6 |
| RP5 NMJ muscle 6/7 | 0.21 | 0.73 | 5 |
| RP2 axon (entry) | 0.98 | 0.02* | 4 |
| RP2 axon (muscle 4) | 0.50 | 0.32 | 6 |
| RP5 axon (entry) | 0.23 | 0.77 | 4 |

### *Rtnl1*^*-*^ *ReepA*^*-*^ *ReepB*^*-*^ mutants display an increased ER lumenal evoked response compared to WT

*Rtnl1*^*-*^ *ReepA*^*-*^ *ReepB*^*-*^ mutant larvae have a disrupted axonal ER network, with larger and fewer tubules, and occasional fragmentation (Yalçın et al. 2017). To test whether these ER changes had functional consequences, we used *Dpr-GMR94G06-GAL4* to examine the cytoplasmic and ER lumen evoked responses at muscle 1 NMJ in these mutants. Mutant NMJs showed an increased evoked response compared to WT in the ER lumen, but not in the cytoplasm (Fig. 4A-B). There was no significant difference in lumen resting fluorescence between mutant and WT (Fig. 4C).

**Fig 4.**
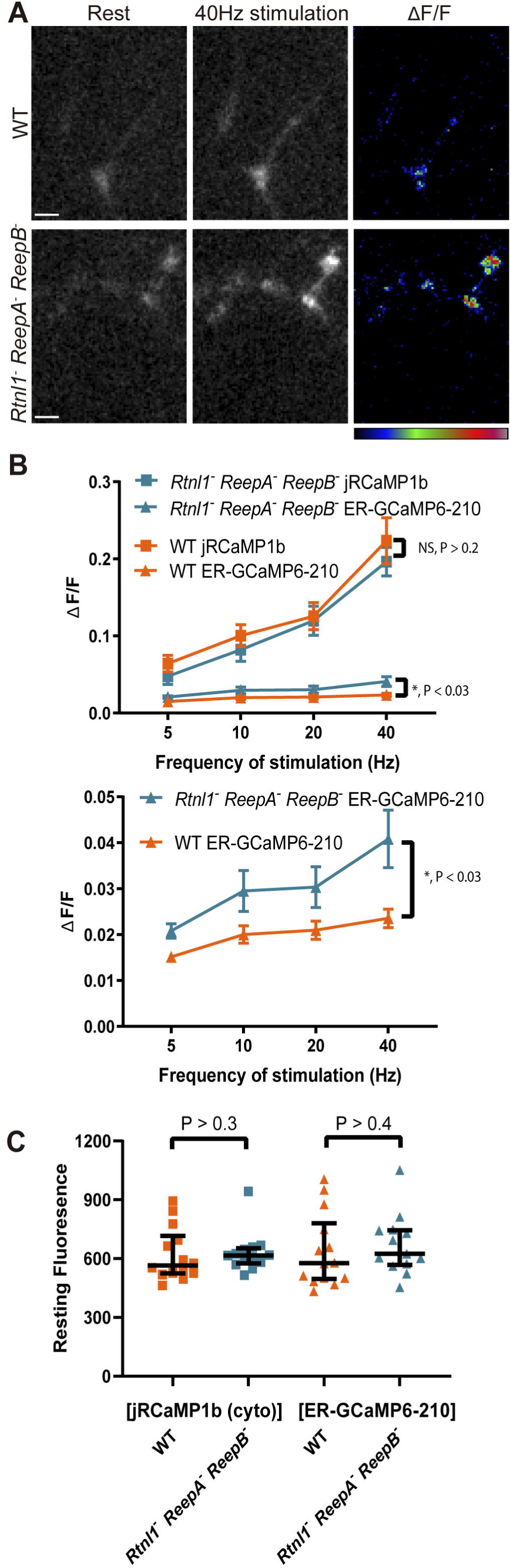
ER lumenal, but not cytoplasmic responses, were significantly increased in *Rtnl1*^*-*^ *ReepA*^*-*^ *ReepB*^*-*^ mutant Type Ib boutons compared to *WT*, with no difference in resting fluorescence intensity. (**A**) Lumenal ER-GCaMP6-210 expressed in aCC using *Dpr-GMR94G06-GAL4*, showed a larger evoked response in Type Ib NMJs at muscle 1 in mutants, compared to *WT*. 40 Hz stimulation panel shows the maximum response of the given GECI following stimulation. GECIs were co-expressed in the same neuron. Pseudocolour bar (for ΔF/F panels) shows low to high ΔF/F; scale bar, 5 µm. (**B**) There was no significant difference between mutant (N=13) and WT (N=14) in cytoplasmic responses (jRCaMP1b) at these boutons stimulated over a range of frequencies, but there was a significant increase in the ER lumenal response (2-way ANOVA). The lower graph shows the ER lumenal response only, with the y-axis expanded. (**C**) There was no significant effect of the triple mutant on the resting fluorescence intensity (from the 40 Hz recording) of either sensor (Mann-Whitney U test).

There was no correlation between cytoplasmic and ER lumenal evoked responses in either mutant or WT at 40 Hz (Fig. S3), supporting the finding in Fig. 3 of a more complex intracellular transfer of calcium. It has been reported that ER lumen [Ca^2+^] affects presynaptic cytoplasmic Ca^2+^ flux (de Juan-Sanz et al. 2017); however we found no correlation here between resting ER fluorescence and peak cytoplasmic fluorescence, in either WT or mutant stimulated at 40 Hz (Fig. S3). The time-to-peak in the cytoplasm or ER lumen was not significantly different between mutant and WT at 40 Hz (Fig. S4). The time for decay to half-maximal in the cytoplasm was also not significantly different between mutant and WT at 40 Hz (Fig. S4). This parameter was not measured in the ER lumen due to the slow temporal dynamics of this flux.

These data suggest a significant increase in ER lumenal evoked responses in *Rtnl1*^*-*^ *ReepA*^*-*^ *ReepB*^*-*^ mutants compared to WT, with no change in ER lumenal resting fluorescence levels, no corresponding change in the cytoplasmic evoked response, and no change in the temporal dynamics of the response.

## Discussion

Direct measurement of ER lumenal Ca^2+^ dynamics is essential for reporting on the physiological and pathophysiological responses of the ER network. An ER lumenal *UAS-GECI*, ER-GCaMP6-210, provided the most robust responses in *Drosophila* motor neurons (Fig. 2), in keeping with results in mammalian hippocampal neurons (de Juan-Sanz et al. 2017). This sensor was also recently successfully cloned and expressed in *Drosophila*, and used to measure Ca^2+^ efflux from the ER in mushroom body Kenyon cells (Handler et al. 2019).

In mutant *Drosophila* lacking various ER-shaping HSP protein homologs, we previously found decreased ER levels compared to WT, or occasional discontinuity of the ER network (Yalçın et al. 2017). As sustained discontinuity could potentially disrupt long-range ER communication along the axon, we wanted to investigate if we could record Ca^2+^ fluxes in WT axons. We analyzed both cytoplasmic and ER lumenal Ca^2+^ fluxes, at three axonal recording positions across two neurons. The amplitude of the ER lumenal evoked response was similar in axons and at the NMJ; however cytoplasmic evoked responses were smaller in axons compared to the NMJ. Even though Ca^2+^ would be fluxing across the ER membrane in response to stimulation, it appears that the two signals do not influence each other in most cases (Fig. 3).

While we observe Ca^2+^ signals along the length of motor axons, we do not find any indication of propagation via a Ca^2+^ wave. Propagation speeds for Ca^2+^ waves in dendrites have been measured at 60-90 µm/s (Nakamura et al. 2002), with a rapid lumenal Ca^2+^ diffusion through dendritic ER at ∼30 µm/s (Choi et al. 2006). Here we find no evidence for any spatial differences in the time to peak fluorescence between different axon positions and the NMJ, both for ER lumen and cytoplasmic evoked responses (Fig. 3); in a typical video frame width of 100 µm (e.g. Fig. S1, S2) we should easily detect propagation at these speeds with our frame rate of 10 Hz. The instantaneous spread of Ca^2+^ responses that we observe is more consistent with them being a consequence of the action potentials generated by stimulation, which propagate so fast that we cannot resolve their spread using our optical recording. Detection of Ca^2+^ waves that propagate along the ER membrane, or through the lumen would probably require recording of ER Ca^2+^ fluxes in the absence of action potentials.

Although the cytoplasmic evoked responses of *Rtnl1*^*-*^ *ReepA*^*-*^ *ReepB*^*-*^ mutants were normal, the ER lumenal evoked responses were significantly larger than WT (Fig. 4). This did not appear to be due to increased resting ER lumenal [Ca^2+^] since resting fluorescence levels showed no significant difference between genotypes, for either ER or cytoplasmic sensors.

In the simplest interpretation, the presynaptic ER lumen appears to take up more Ca^2+^ on stimulation in the *Rtnl1*^*-*^ *ReepA*^*-*^ *ReepB*^*-*^ mutant compared to WT. Since cytoplasmic Ca^2+^ responses are unaffected in mutants, increased cytosolic Ca^2+^ cannot be an explanation for the increased ER response in mutants, implying that the increased ER response is due to intrinsic changes in Ca^2+^-handling properties of the mutant ER, consistent with the roles of reticulon and REEP proteins in directly shaping ER architecture. Furthermore, increased uptake of Ca^2+^ by mutant ER does not lead to decreased cytoplasmic [Ca^2+^]; however, we might not detect very local reductions in [Ca^2+^] close to the ER, if masked by excess cytoplasmic fluorescence away from its immediate vicinity. Increasing bath [Ca^2+^] can rescue transmitter release phenotypes in *atl* and *Rtnl1* mutants (Summerville et al. 2016); reducing bath [Ca^2+^] in our setup might reveal cytoplasmic dysfunction not apparent at higher bath [Ca^2+^].

Could the increased ER lumenal Ca^2+^ responses in mutants have any effects on cell physiology, if there is no corresponding increase in [Ca^2+^] in the cytoplasm, where many Ca^2+^ effectors reside? There are Ca^2+^ effectors in the ER lumen such as calreticulin (Opas et al. 1991) although the protein-folding chaperone role of calreticulin is more likely to place its main function in rough rather than smooth ER. However, altered lumenal Ca^2+^ responses could also affect physiology of organelles that obtain Ca^2+^ from the ER, including mitochondria and endo/lysosomal organelles (Raffaello et al. 2016)]. These effects could endure well beyond the original train of action potentials, due to the slow kinetics of Ca^2+^ uptake and release by the ER, with lumenal [Ca^2+^] not always returning to baseline during our recordings. Furthermore aberrant ER Ca^2+^ handling has been postulated as a trigger for neurodegeneration, so what we are observing may well reflect deleterious changes in cellular health (Stutzmann and Mattson 2011). In support of our findings, mutation or knockdown of genes with a role in ER shaping, including *atl, Rtnl1* and *VAP*, also affect aspects of presynaptic function including neurotransmitter release and synaptic vesicle density, and the cytoplasmic presynaptic Ca^2+^ response (Summerville et al. 2016, De Gregorio et al. 2017, Lindhout et al. 2019). Taken together, this data strongly suggests a role for ER in modulating presynaptic function, and highlights the importance of having an intact, functional ER network.

In conclusion, we have demonstrated the use of ER lumenally targeted GECIs for use in *Drosophila*, successfully recording fluxes at both the NMJ and in the axon. We have demonstrated the utility of these sensors in identifying functional differences between WT and *Rtnl1*^*-*^ *ReepA*^*-*^ *ReepB*^*-*^ mutant flies, whose axonal ER network it is known to be affected. We anticipate these will be useful additions to a *Drosophila* Ca^2+^ imaging toolkit in the future, to further our understanding of HSP, and neurodegeneration more broadly.

## Author Contributions

MKO performed all the cloning and Ca^2+^ imaging experiments, and together with CJO’K designed the experiments. JJPM performed the confocal imaging acquisition. JO’S performed preliminary experiments and contributed to data analysis. TW provided initial training and support for Ca^2+^ imaging experiments, and assisted in experimental design. The paper was written by MKO and CJO’K with comments and edits by JJPM, TJW and JO’S.

## Acknowledgments

We thank The University of Cambridge Department of Genetics Fly Facility for providing the *Drosophila* microinjection service, Timothy Ryan for providing the FCK(1.3)GW-ER-GCaMP6-210 plasmid and Bloomington *Drosophila* Stock Centre for stocks.

