## Supplementary Material for "Endoplasmic reticulum (ER) lumenal indicators in *Drosophila* reveal effects of HSP-related mutations on ER calcium dynamics"

### 1 Supplementary Figures and Tables

#### 1.1 Supplementary Figures

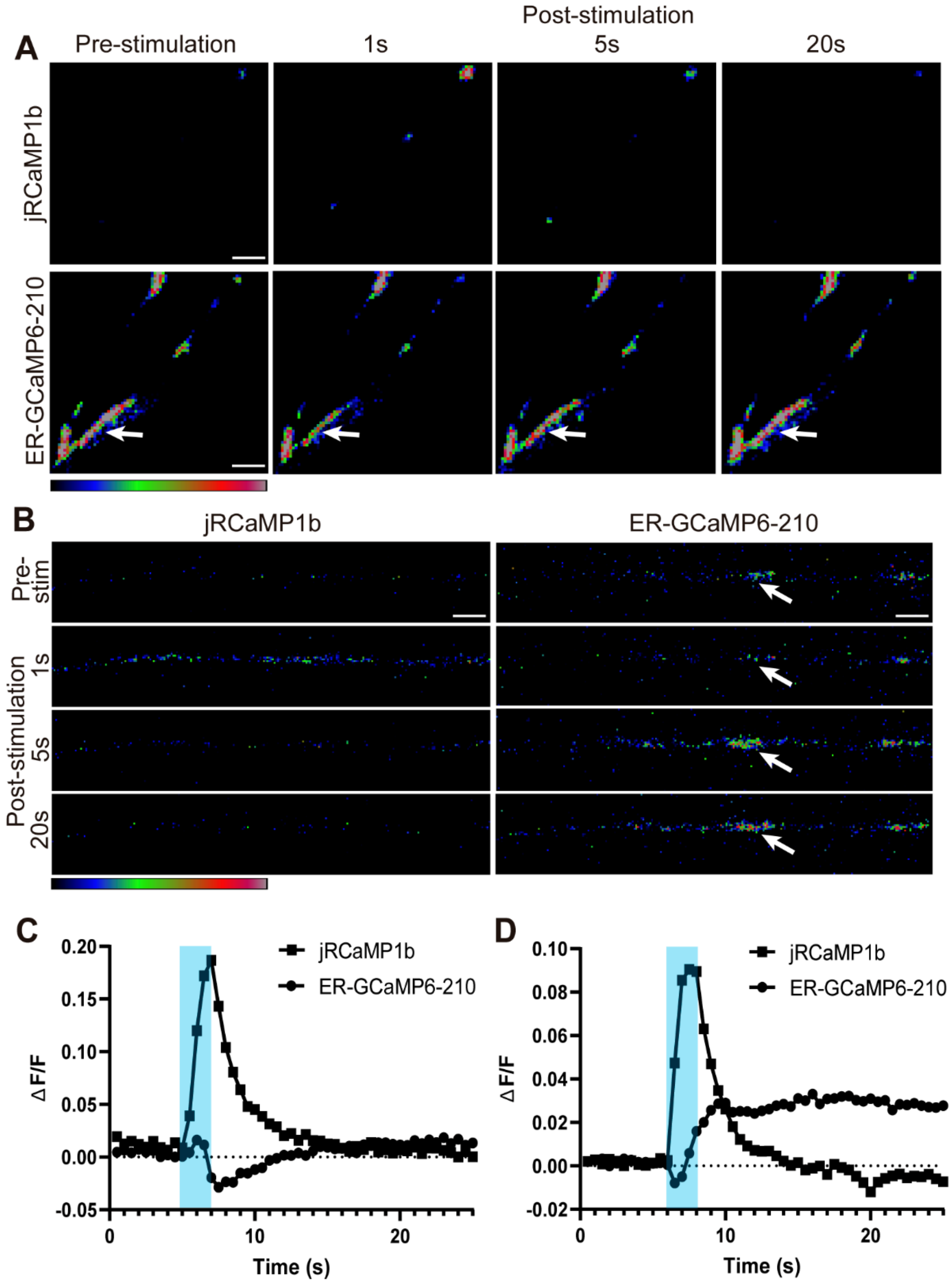


**Supplementary Figure 1.** Examples of ER lumenal response to electrical stimulation, showing a small sharp decrease in ER lumenal fluorescence immediately following stimulation. Such a decrease occurred in approximately 10% of nerves. (**A**) Representative pseudocolour images of responses of ER-GCaMP6-210 and jRCaMP1b, expressed in the aCC Type Ib neuron using *Dpr-GMR94G06-GAL4*, to 20 Hz stimulation at NMJ muscle 1. In this example the decrease in ER fluorescence (change in pseudocolor from gray to yellows and blues, shown by arrows) is followed by a return to baseline fluorescence. Pseudocolour bar shows low to high relative intensity; scale bar, 5 µm. (**B**) Representative pseudocolour images of responses of ER-GCaMP6-210 and jRCaMP1b, expressed in the RP2 Type Is neuron using *FMR-GMR27E09-GAL4*, to 40 Hz stimulation in the axon (muscle 4). In this example a localized decrease in ER fluorescence is followed by a slow increase in fluorescence, above baseline levels (arrows). Pseudocolour bar shows low to high relative intensity; scale bar, 5 µm. (**C**) Graphical representation of response in (**A**), shading represents the stimulation period. (**D**) Graphical representation of response in (**B**), shading represents the stimulation period.

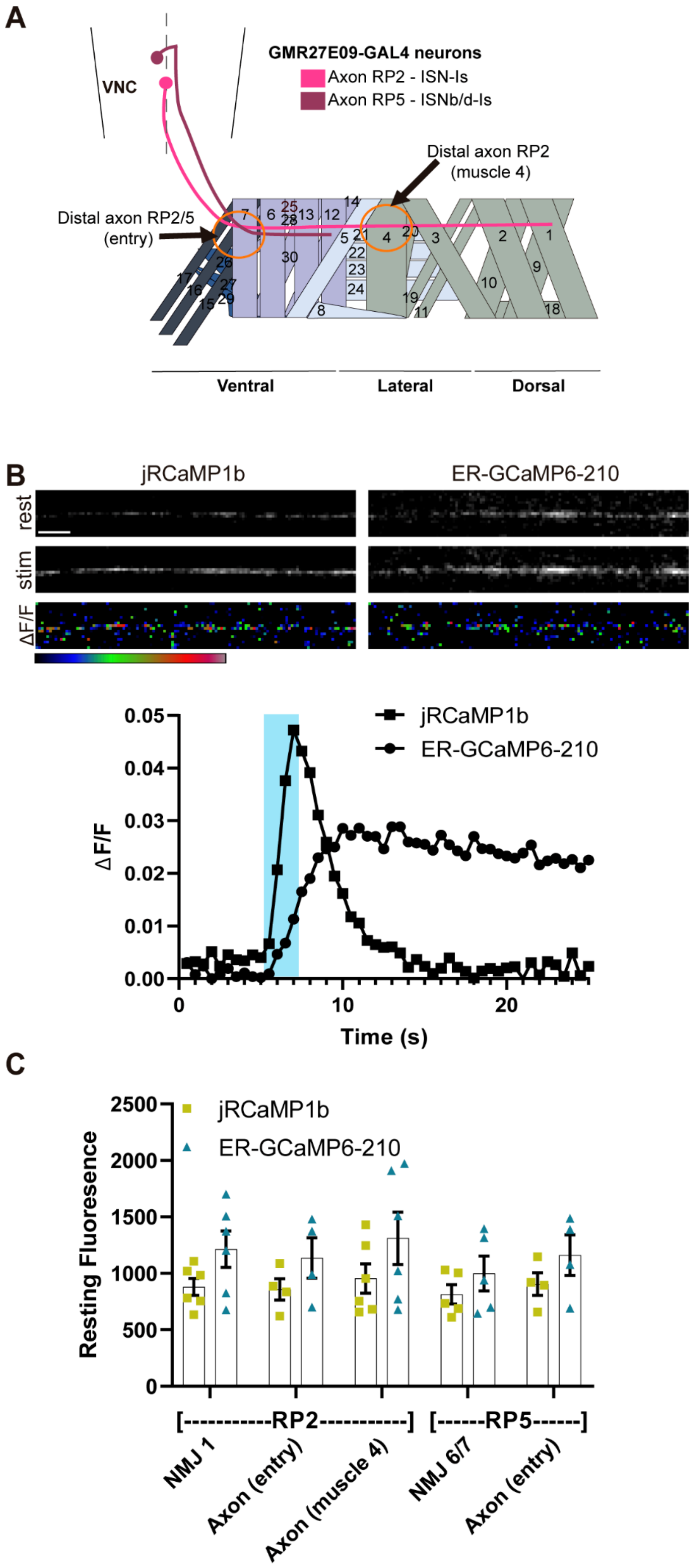
A
VNC
GMR27E09-GAL4 neurons
Axon RP2 - ISN-Is
Axon RP5 - ISNb/d-Is
Distal axon RP2
(muscle 4)
Distal axon RP2/5
(entry)
Ventral
Lateral
Dorsal
B
jRCaMP1b
ER-GCaMP6-210
rest
stim
ΔF/F
ΔF/F
Time (s)
C
jRCaMP1b
ER-GCaMP6-210
Resting Fluoresence
[-----------RP2-----------]
[------RP5------]
NMJ 1
Axon (entry)
Axon (muscle 4)
NMJ 6/7
Axon (entry)

**Supplementary Figure 2.** Locations of axonal recordings from neurons RP2 and RP5, labeled using *FMR-GMR27E09-GAL4*. (**A**) Scheme of innervation of an abdominal hemisegment highlighting Type Is motoneurons that express *FMR-GMR27E09-GAL4*. Axonal recording points are circled. Adapted from Perez-Moreno and O'Kane (2019). (**B**) Representative grayscale images of jRCaMP1b and ER-GCaMP6-210 responses to 20 Hz stimulation at axon RP2 (over muscle 4). Stim panel shows the maximum response of the given GECI following stimulation. GECIs were co-expressed in the same axon. Scale bar, 5 µm. Pseudocolour panels show ΔF/F, with bar below showing low to high flux. Graph shows timecourse of ΔF/F for a typical axonal response. **(C)** Resting fluorescence intensity (from the 40 Hz recording) was consistent across axonal and NMJ locations.

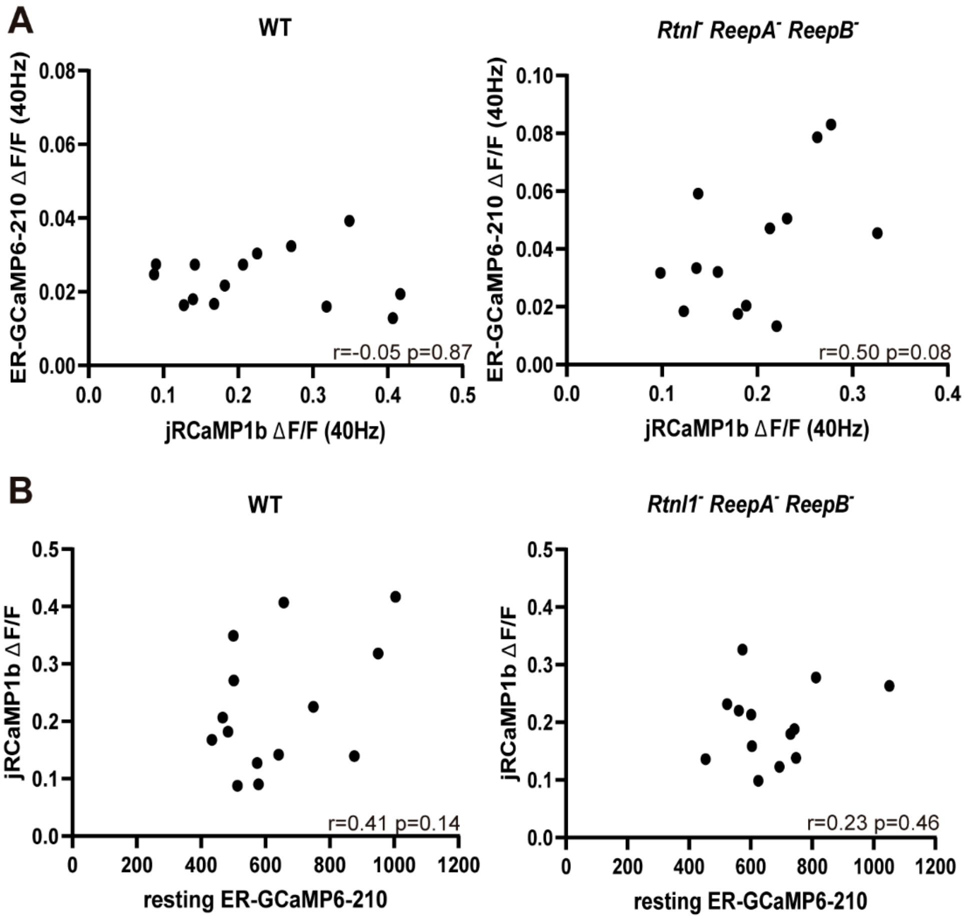


**Supplementary Figure 3.** Measuring correlations between ER lumenal ER-GCaMP6-210 and cytosolic jRCaMP1b $Ca^{2+}$ responses. There was no correlation (Pearson's correlation coefficient) found at muscle 1 NMJ in the aCC Type Ib neuron using *Dpr-GMR94G06-GAL4* (**A**) between the cytoplasmic and ER lumenal evoked responses in either WT (N=14) or *Rtnl1⁻ ReepA⁻ ReepB⁻* mutant (N=13) at 40 Hz, or (**B**) between resting ER fluorescence and peak cytoplasmic fluorescence at 40 Hz in either WT (N=14) or *Rtnl1⁻ ReepA⁻ ReepB⁻* mutant (N=13).

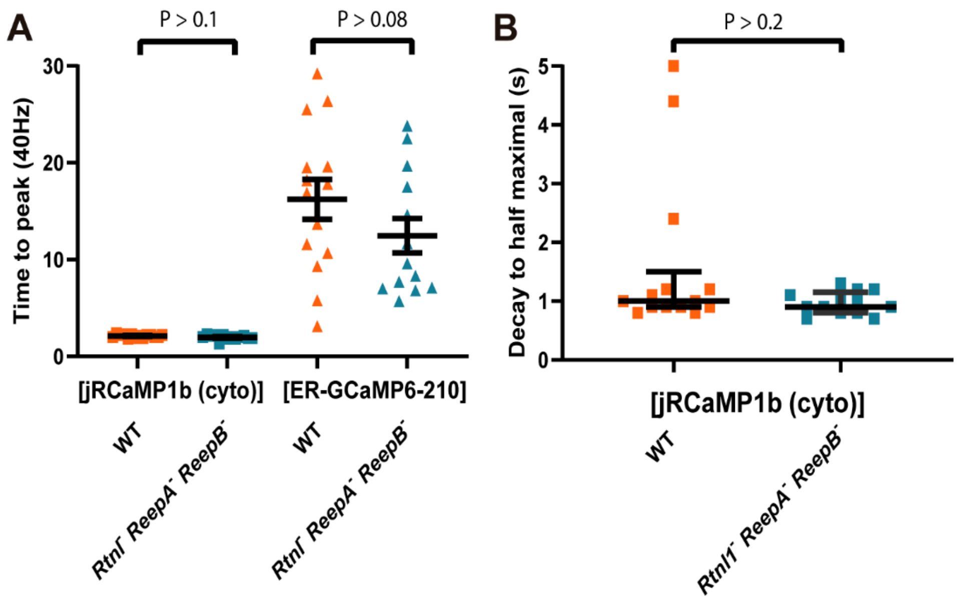


**Supplementary Figure 4.** Measuring the temporal dynamics of the ER lumenal ER-GCaMP6-210 and cytosolic jRCaMP1b $Ca^{2+}$ responses. Using *Dpr-GMR94G06-GAL4* at muscle 1 NMJ in the aCC Type Ib neuron, (**A**) the time-to-peak in the cytoplasm or ER lumen (Student's t-test) was not significantly different between WT (N=14) and *Rtnl1*⁻ *ReepA*⁻ *ReepB*⁻ mutant (N=13) at 40Hz. Error bars show SEM. (**B**) Time for decay to half maximal in the cytoplasm was also not significantly different between WT (N=14) and *Rtnl1*⁻ *ReepA*⁻ *ReepB*⁻ mutant (N=13) at 40Hz (Mann-Whitney U test). Data show median and interquartile range.

**Supplementary File 1.** Cloning scheme for generating *p17xUASTattB*, *p17xUASTattB-ER-GCaMP6-210 p17xUASTattB-CEPIA3-ER*, and *p17xUASTattB-CEPIA4-ER*.

*p17xUASTattB* was created by using PCR to amplify the 20xUAS site region of *pJFRC-20xUAS-Gateway modified (ccdB –ve)*, including the existing *Hin*dIII site in the backbone of this vector, and adding a second *Hin*dIII site to the tail end of the reverse primer (tail region underlined in schematic) (Slide 1). This fragment, and the *pUASTattB-5xUAS* Gateway-compatible vector were both digested with *Hin*dIII enzyme, linearizing the *pUASTattB-5xUAS* Gateway-compatible vector (Slide 2). Upon ligation of the fragment and the vector, a Gateway-compatible vector containing the ccdB gene and 17 UAS sites was obtained (Slide 3).

*p17xUASTattB-ER-GCaMP6-210* was created by using PCR to amplify the calreticulin signal peptide, sensor and KDEL retention sequence from *ER-GCaMP6-210* (Table 1). Tails (underlined in schematic) were added to the primers to add attB1 and Kozak sequence to the 5’ end of the forward primer, and attB2 to the 5’ end of the reverse primer. Colored and numbered amino acids correspond to coding regions of the plasmids, corresponding to the colors in the diagram (Slide 4). Gateway$^{TM}$ cloning was then used to recombine the fragments into *p17xUASTattB* (Slide 5,6).

*p17-UASTattB-CEPIA3-ER* and *p17-UASTattB-CEPIA4-ER* were created by using PCR to amplify the sensor and myc tag region from *pCMV CEPIA3mt* and *pCMV CEPIA4mt* (Table 1), excluding the mitochondrial signal sequence. Tails (underlined in schematic) were added to the primers to add attB1, Kozak and BiP signal sequence to the 5’ end of the forward primer, and attB2, stop codon and HDEL retention sequence to the 5’ end of the reverse primer. Colored and numbered amino acids correspond to coding regions of the plasmids, corresponding to the colors in the diagram (Slide 7). Gateway$^{TM}$ cloning was then used to recombine the fragments into *p17xUASTattB* (Slide 8,9).

**Supplementary File 2.** R scripts for $Ca^{2+}$ flux analysis. The first file was used for analysis of each individual recording, and the combined analysis was used for each subgroup to calculate average values. Annotations are in scripts.

**Supplementary Video 1.** Representative presynaptic responses to 20 Hz stimulation, of ER-GCaMP6-210 in ER lumen (green) and jRCaMP1b in cytoplasm (magenta) at muscle 6/7 NMJ, driven by *FMR-GMR27E09-GAL4* in the RP5 Type Is neuron. ROI outline in yellow. Scale bar 5 µm.

**Supplementary Video 2.** Representative axonal $Ca^{2+}$ responses to 20 Hz stimulation, of ER-GCaMP6-210 in ER lumen (green) and jRCaMP1b in cytoplasm (magenta) at axon of neuron RP2 Type Is neuron (over muscle 4), driven by *FMR-GMR27E09-GAL4*. Scale bar 5 µm.
