## Supplementary File 1 for "Endoplasmic reticulum (ER) lumenal indicators in *Drosophila* reveal effects of HSP-related mutations on ER calcium dynamics"

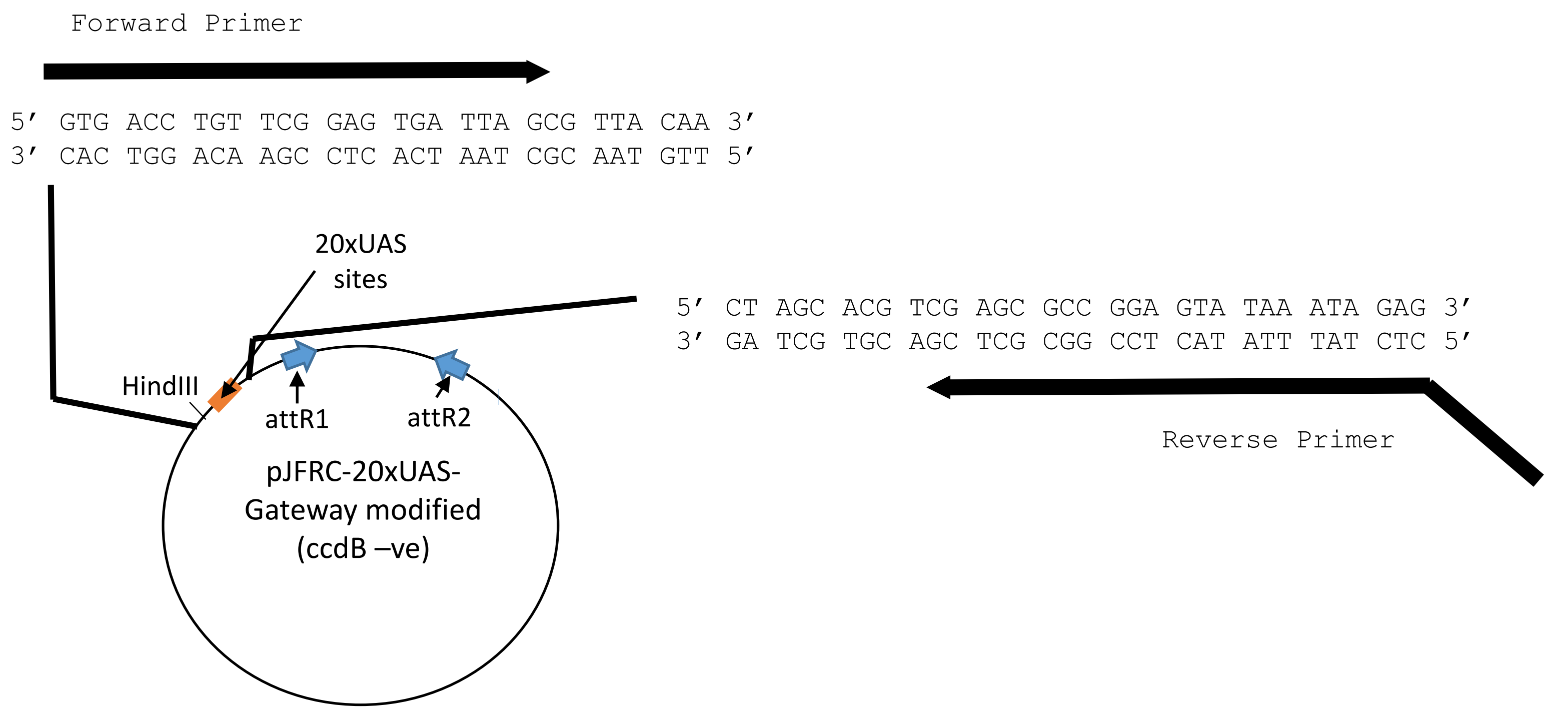


Forward Primer:

5′ GTG ACC TGT TCG GAG TGA TTA GCG 3′

Reverse Primer:

5′ AAG CTT CTC TAT TTA TAC TCC GGC GCT CG 3′
**HindIII**

PCR ↓

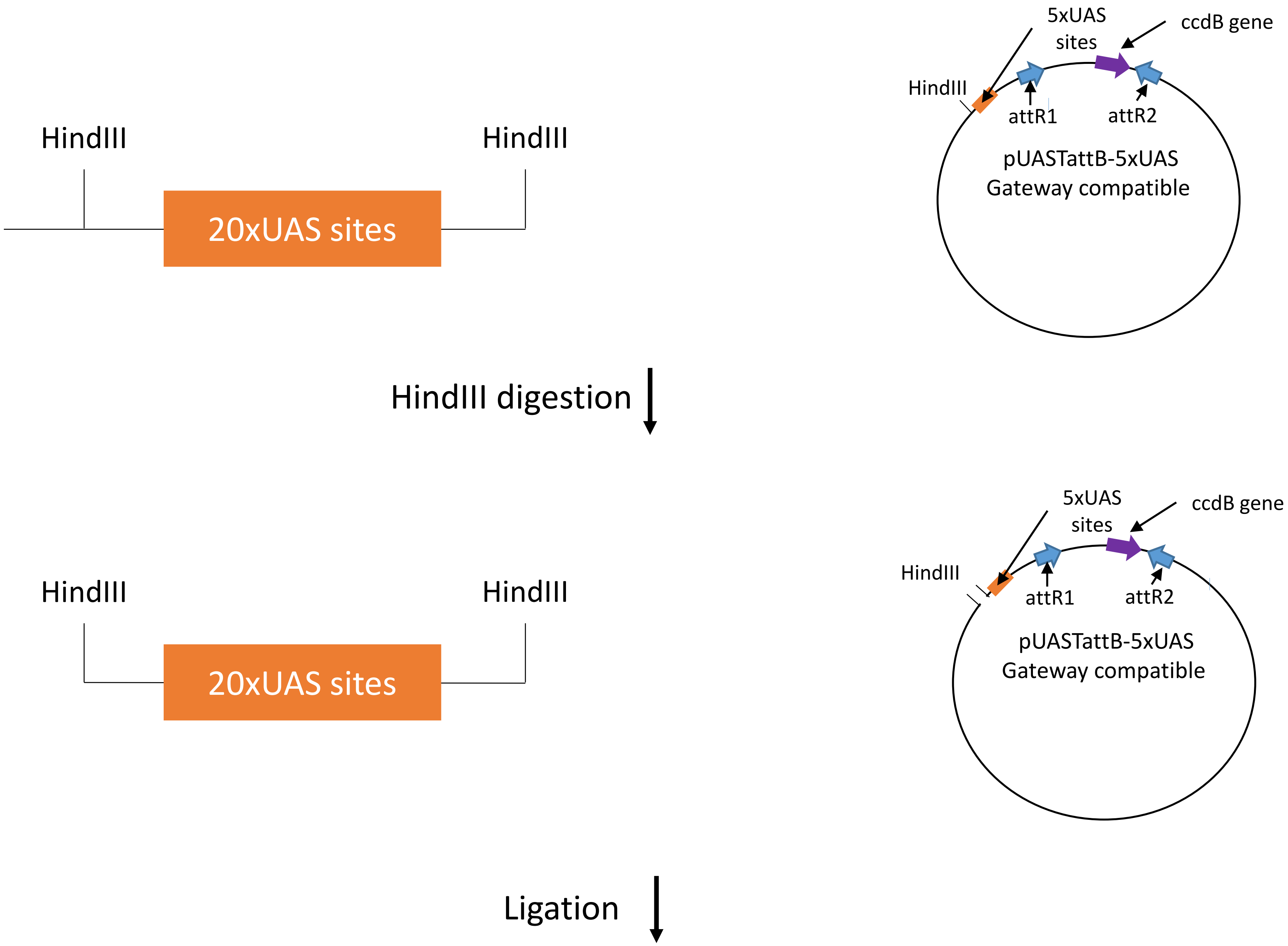

5xUAS sites
ccdB gene
HindIII
attR1
attR2
pUASTattB-5xUAS
Gateway compatible
HindIII
HindIII
20xUAS sites
HindIII digestion
5xUAS sites
ccdB gene
HindIII
attR1
attR2
pUASTattB-5xUAS
Gateway compatible
HindIII
HindIII
20xUAS sites
Ligation

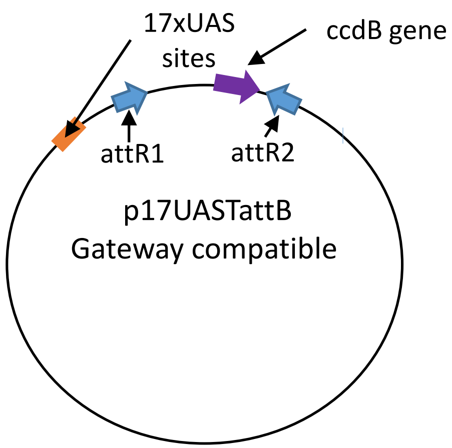
17xUAS
sites
ccdB gene
attR1
attR2
p17UASTattB
Gateway compatible

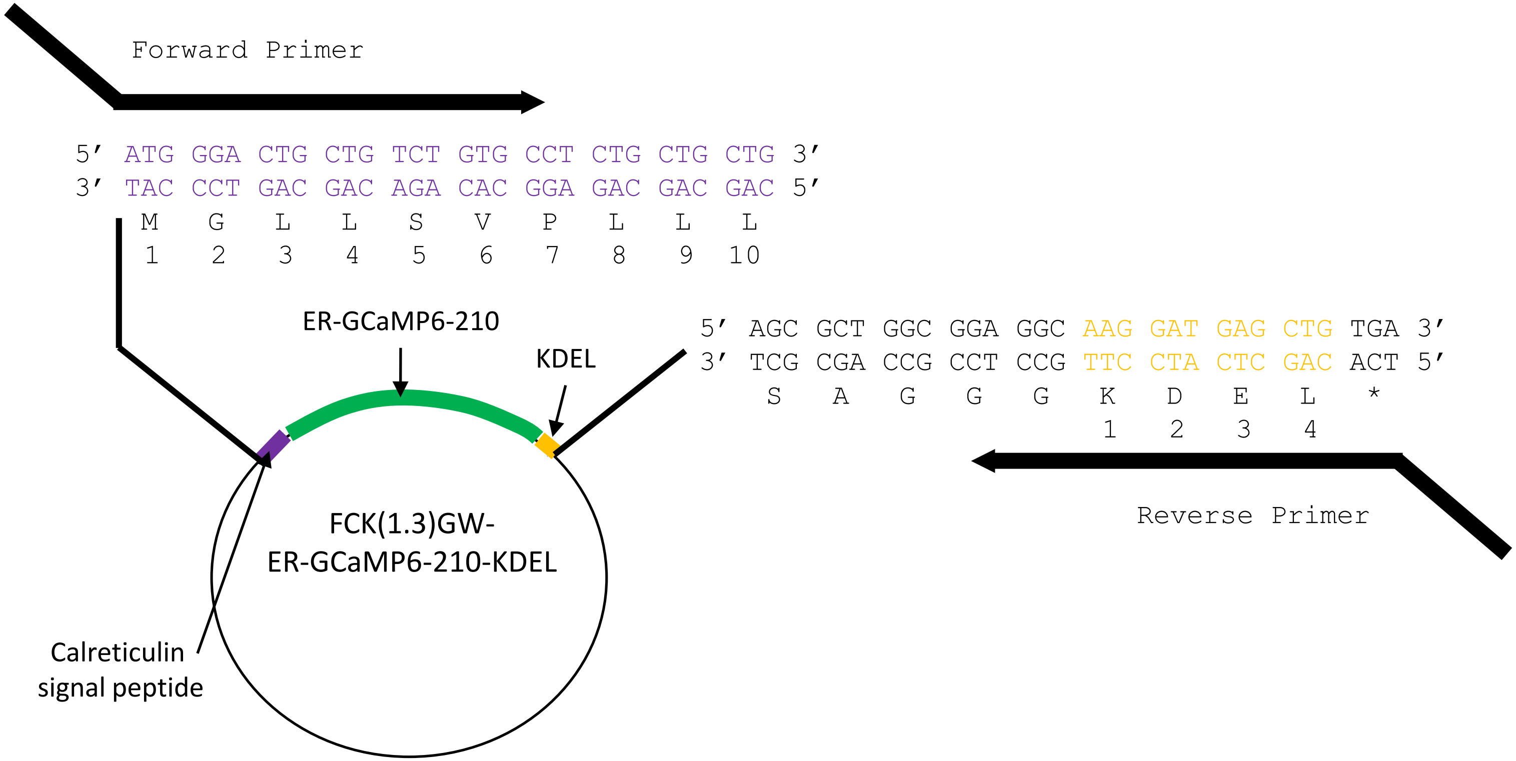


Forward Primer:

5′ GGGG ACA AGT TTG TAC AAA AAA GCA GGC TTA CAAAC ATG GGA CTG CTG TCT GTG C 3′

**attB1** **Kozak** M G L L S V
1 2 3 4 5 6

Reverse Primer:

5′ GGGG AC CAC TTT GTA CAA GAA AC TGG GTT TCA CAG CTC ATC CTT GCC T 3′

**attB2** * L E D K G
4 3 2 1

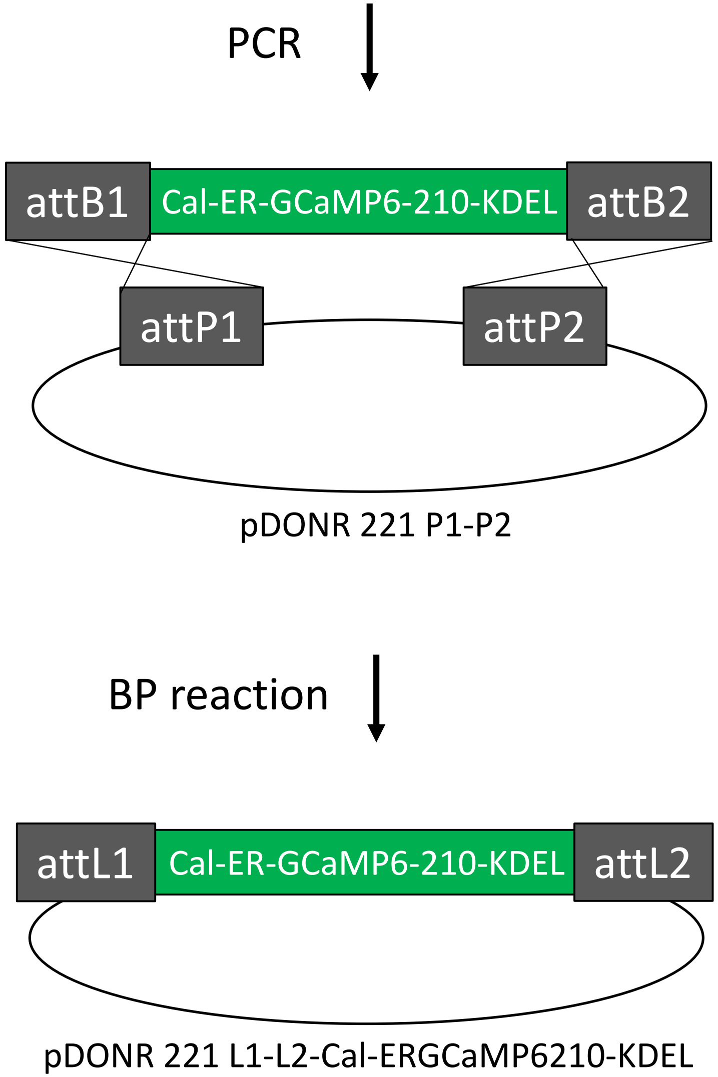
PCR
attB1
Cal-ER-GCaMP6-210-KDEL
attB2
attP1
attP2
pDONR 221 P1-P2
BP reaction
attL1
Cal-ER-GCaMP6-210-KDEL
attL2
pDONR 221 L1-L2-Cal-ERGCaMP6210-KDEL

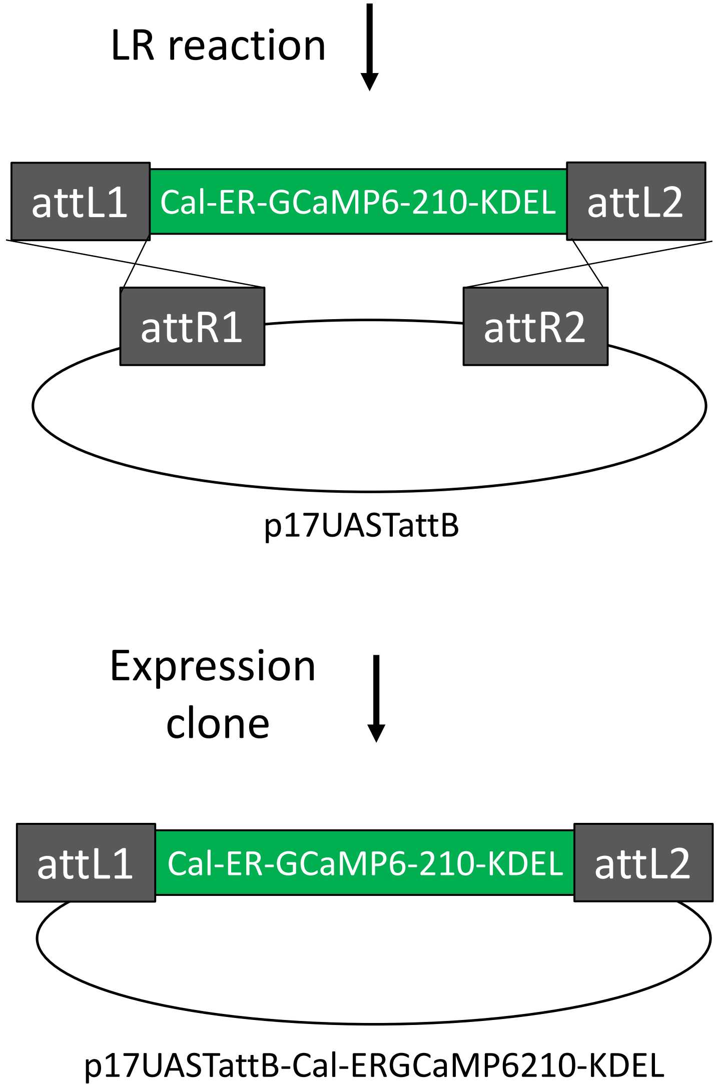
LR reaction
attL1
Cal-ER-GCaMP6-210-KDEL
attL2
attR1
attR2
p17UASTattB
Expression
clone
attL1
Cal-ER-GCaMP6-210-KDEL
attL2
p17UASTattB-Cal-ERGCaMP6210-KDEL

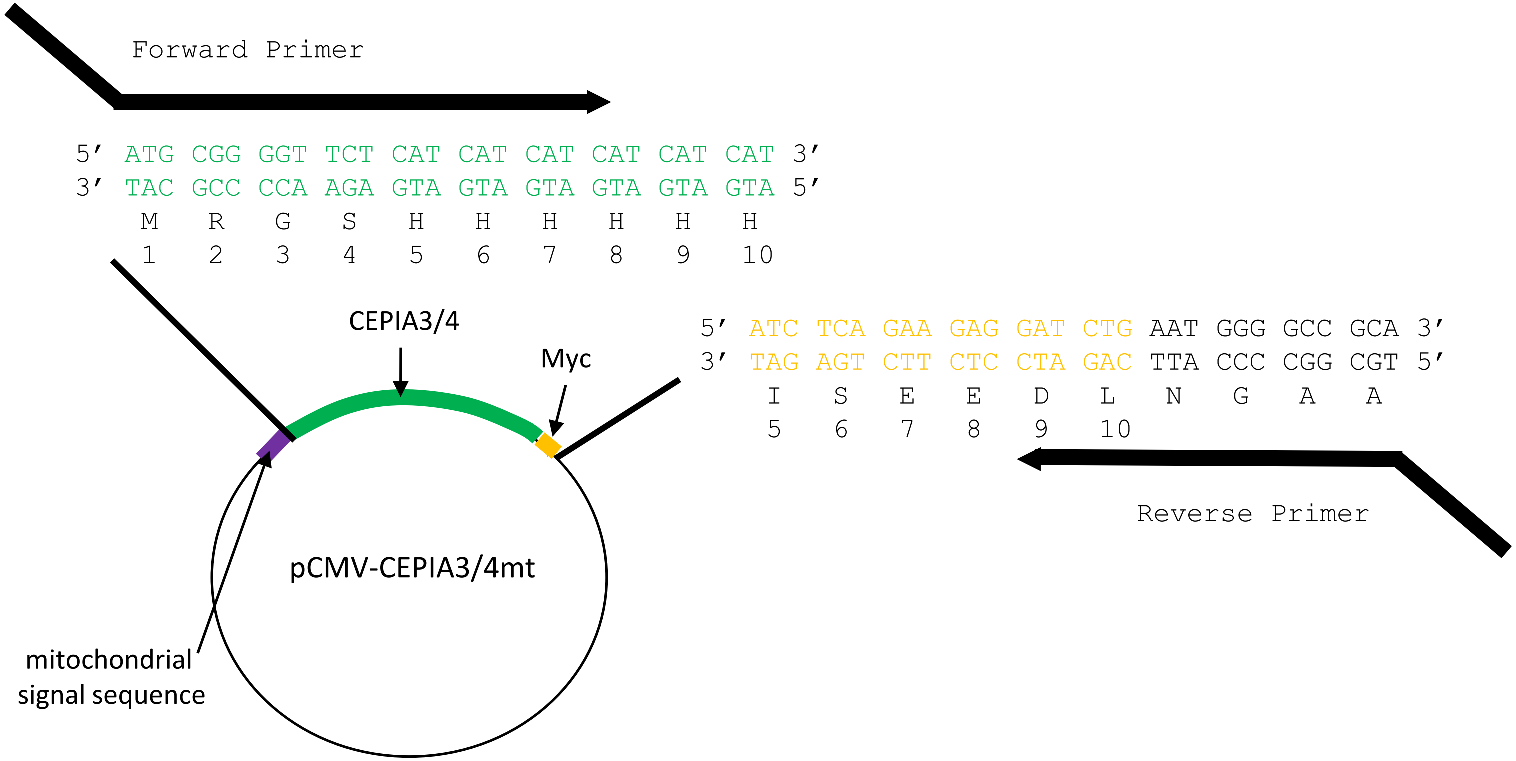


Forward Primer:

5′ GGGG ACA AGT TTG TAC AAA AAA GCA GGC TTA CAAAC ATG AAG TTA TGC ATA TTA CTG GCC GTC GTG GCC TTT GTT GGC CTC

attB1 | Kozak | M K L C I L L A V V A F V G L

BiP signal sequence

TCG CTC GGG ACC GGT GGC GGA ATG CGG GGT TCT CAT CAT CAT C 3′

S L G T G G G M R G S H H H

1 2 3 4 5 6 7

Reverse Primer:

5′ GGGG AC CAC TTT GTA CAA GAA AC TGG GTT TCA CAA TTC GTC GTG TGC GGC CCC ATT CAG ATC 3′

attB2 | * L E D H A A G N L D

HDEL | 10 9

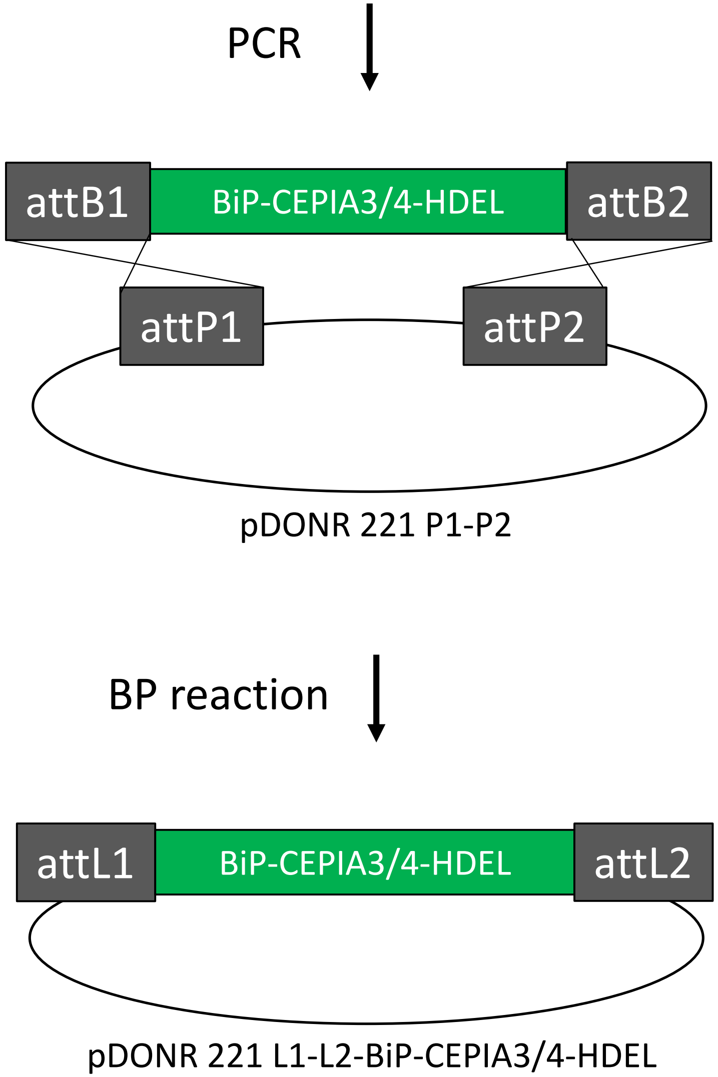
PCR
attB1
BiP-CEPIA3/4-HDEL
attB2
attP1
attP2
pDONR 221 P1-P2
BP reaction
attL1
BiP-CEPIA3/4-HDEL
attL2
pDONR 221 L1-L2-BiP-CEPIA3/4-HDEL

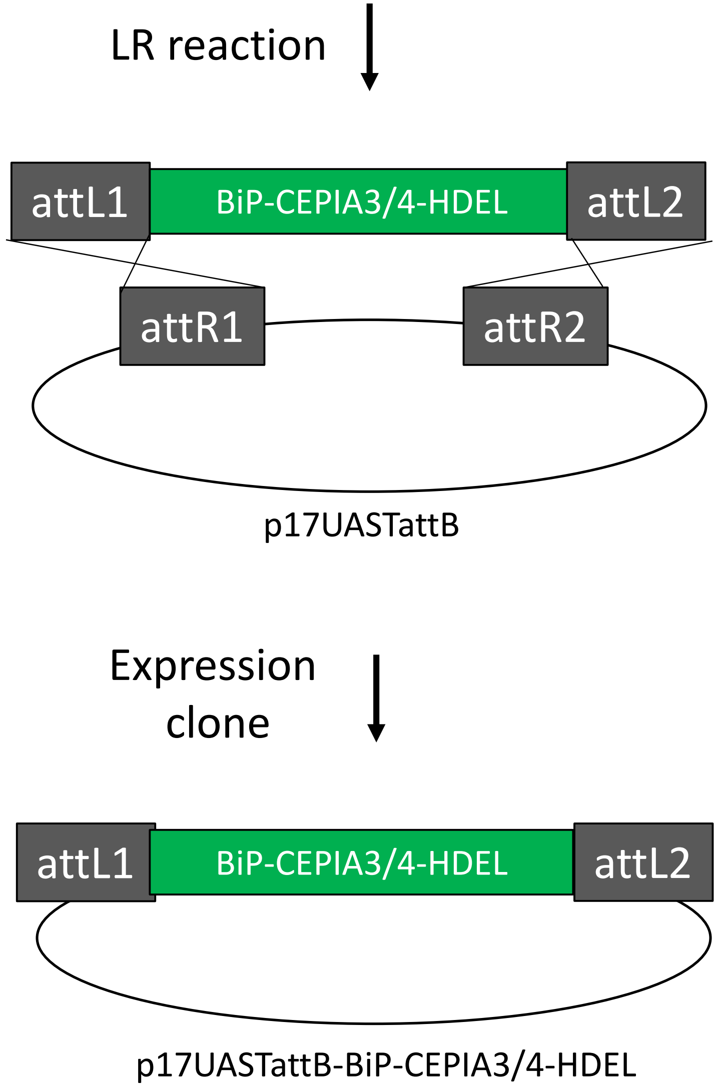
LR reaction
attL1
BiP-CEPIA3/4-HDEL
attL2
attR1
attR2
p17UASTattB
Expression
clone
attL1
BiP-CEPIA3/4-HDEL
attL2
p17UASTattB-BiP-CEPIA3/4-HDEL
