## Supplementary File 2 for "Endoplasmic reticulum (ER) lumenal indicators in *Drosophila* reveal effects of HSP-related mutations on ER calcium dynamics"

Figure 2 3 4 individual analysis R script

```{r freq analysis jRCaMP1b}
#note that this script can be used for all individual analysis in Fig2,3 and 4 by changing the files names to reflect different data sets and copying the code to include extra frequencies.

#note only the 5Hz code is annotated. This is repeated for each frequency, with the number at the end of f_main, zeroed_main, time, peak and timetohalf denoting the frequency at which this values come from

#read in individual txt files that are created when using time series analyser plugin in Fiji. ROI was always named "main"

#5Hz file
filename_main <- "prep1_nerve1_muscle_5Hz_red.txt"
intensity_main <- read.table(filename_main, header=TRUE)
#f_main5 is the averaged background fluorescent intensity - from frame 50-80
f_main5 <- mean(intensity_main$main[50:80])
f_main5
#deltaf_main is df/f
deltaf_main <- (intensity_main$main-f_main5)/f_main5
#find the minimum df/f value to zero the graph
min_main <- which.min(deltaf_main)
#zeroed_main5 is the zeroed data
zeroed_main5 <- deltaf_main
for (i in 1:length(deltaf_main)){
  zeroed_main5[i] <- deltaf_main[i]-deltaf_main[min_main]
}

#time to peak was looked for between frames 110-140
timetopeak_period <- zeroed_main5[110:140]
max_main <- which.max(timetopeak_period)
#peak is the maximum df/f
peak5 <- timetopeak_period[max_main]
#convert to seconds
time5 <- (10+max_main)*0.1

#half maximal recovery time was searched for between 120-200 frames
half_max <- peak5/2
half_max_period <- zeroed_main5[120:200]
half_timepoint <- which.min(abs(half_max_period - half_max))
half <- half_max_period[half_timepoint]
#convert to seconds - the actual time
half_time <- (20+half_timepoint)*0.1
#the time to half maximal recovery
timetohalf5 <- half_time-time5

filename_main <- "prep1_nerve1_muscle_10Hz_red.txt"
intensity_main <- read.table(filename_main, header=TRUE)
f_main10 <- mean(intensity_main$main[50:80])
f_main10
deltaf_main <- (intensity_main$main-f_main10)/f_main10
min_main <- which.min(deltaf_main)
zeroed_main10 <- deltaf_main
for (i in 1:length(deltaf_main)){
 zeroed_main10[i] <- deltaf_main[i]-deltaf_main[min_main]
}

timetopeak_period <- zeroed_main10[110:140]
max_main <- which.max(timetopeak_period)
peak10 <- timetopeak_period[max_main]
time10 <- (10+max_main)*0.1

half_max <- peak10/2
half_max_period <- zeroed_main10[120:200]
half_timepoint <- which.min(abs(half_max_period - half_max))
half <- half_max_period[half_timepoint]
half_time <- (20+half_timepoint)*0.1
timetohalf10 <- half_time-time10

filename_main <- "prep1_nerve1_muscle_20Hz_red.txt"
intensity_main <- read.table(filename_main, header=TRUE)
f_main20 <- mean(intensity_main$main[50:80])
f_main20
deltaf_main <- (intensity_main$main-f_main20)/f_main20
min_main <- which.min(deltaf_main)
zeroed_main20 <- deltaf_main
for (i in 1:length(deltaf_main)){
  zeroed_main20[i] <- deltaf_main[i]-deltaf_main[min_main]
}

timetopeak_period <- zeroed_main20[110:140]
max_main <- which.max(timetopeak_period)
peak20 <- timetopeak_period[max_main]
time20 <- (10+max_main)*0.1

half_max <- peak20/2
half_max_period <- zeroed_main20[120:200]
half_timepoint <- which.min(abs(half_max_period - half_max))
half <- half_max_period[half_timepoint]
half_time <- (20+half_timepoint)*0.1
timetohalf20 <- half_time-time20

filename_main <- "prep1_nerve1_muscle_40Hz_red.txt"
intensity_main <- read.table(filename_main, header=TRUE)
f_main40 <- mean(intensity_main$main[50:80])
f_main40
deltaf_main <- (intensity_main$main-f_main40)/f_main40
min_main <- which.min(deltaf_main)
zeroed_main40 <- deltaf_main
for (i in 1:length(deltaf_main)){
  zeroed_main40[i] <- deltaf_main[i]-deltaf_main[min_main]
}

timetopeak_period <- zeroed_main40[110:140]
max_main <- which.max(timetopeak_period)
peak40 <- timetopeak_period[max_main]
time40 <- (10+max_main)*0.1

half_max <- peak40/2
half_max_period <- zeroed_main40[120:200]
half_timepoint <- which.min(abs(half_max_period - half_max))
half <- half_max_period[half_timepoint]
half_time <- (20+half_timepoint)*0.1
timetohalf40 <- half_time-time40

#put all the zereoed_main data into a matrix and save this to a txt file
zeroed_main_avg <- array(c(zeroed_main5, zeroed_main10, zeroed_main20, zeroed_main40),dim = c(1000,4))
m0 <- c("zeroed_main5", "zeroed_main10", "zeroed_main20", "zeroed_main40")
colnames(zeroed_main_avg) <- m0
write.table(zeroed_main_avg, "190815_prep1_nerve1_muscle1_red.txt")

#put all the f_main values into an array and save this to a txt file
f_main_list <- c(f_main5,f_main10,f_main20, f_main40)
write.table(f_main_list, "190815_prep1_nerve1_muscle1_red_bg.txt")

#put all the time values into an array and save this to a txt file
time_list <- c(time5,time10,time20, time40)
time_list
write.table(time_list, "190815_prep1_nerve1_muscle1_red_time.txt")

#put all the peak values into an array and save this to a txt file
peak_list <- c(peak5,peak10,peak20, peak40)
peak_list
write.table(peak_list, "190815_prep1_nerve1_muscle1_red_peak.txt")

#put all the timetohalf values into an array and save this to a txt file
timetohalf_list <- c(timetohalf5,timetohalf10,timetohalf20, timetohalf40)
timetohalf_list
write.table(timetohalf_list, "190815_prep1_nerve1_muscle1_red_timetohalf.txt")

#these txt files are fed into combined analyses scripts
```

```{r freq analysis ER lumenal}

#the jRCaMP1b code is annotated. The code here is identical except it is for the ER lumenal txt files

filename_main <- "prep1_nerve1_muscle2_5Hz_green.txt"
intensity_main <- read.table(filename_main, header=TRUE)
fmain5_main <- intensity_main$main
f_main5 <- mean(intensity_main$main[50:80])
f_main5
deltaf_main <- (intensity_main$main-f_main5)/f_main5
deltaf_main5 <- (intensity_main$main-f_main5)/f_main5
min_main <- which.min(deltaf_main)
zeroed_main5g <- deltaf_main
for (i in 1:length(deltaf_main)){
  zeroed_main5g[i] <- deltaf_main[i]-deltaf_main[min_main]
}
avged_main5g <- seq(1,80,1)
j=1
for (i in 1:length(avged_main1g)){
  avg <- mean(zeroed_main5g[j:(j+4)])
  avged_main5g[i] <- avg
  j=j+5
}
timetopeak_period <- zeroed_main5g[110:400]
max_main <- which.max(timetopeak_period)
peak <- timetopeak_period[max_main]
time5 <- (10+max_main)*0.1

timetopeak_period <- avged_main5g[24:36]
max_main <- which.max(timetopeak_period)
peak5 <- timetopeak_period[max_main]
#time5 <- (4+max_main)*0.5

half_max <- peak5/2
half_max_period <- avged_main5g[32:80]
half_timepoint <- which.min(abs(half_max_period - half_max))
half <- half_max_period[half_timepoint]
half_time <- (12+half_timepoint)*0.5
timetohalf5 <- half_time-time5

filename_main <- "prep1_nerve1_muscle2_10Hz_green.txt"
intensity_main <- read.table(filename_main, header=TRUE)
fmain10_main <- intensity_main$main
f_main10 <- mean(intensity_main$main[50:80])
f_main10
deltaf_main <- (intensity_main$main-f_main10)/f_main10
deltaf_main10 <- (intensity_main$main-f_main10)/f_main10
min_main <- which.min(deltaf_main)
zeroed_main10g <- deltaf_main
for (i in 1:length(deltaf_main)){
  zeroed_main10g[i] <- deltaf_main[i]-deltaf_main[min_main]
}
avged_main10g <- seq(1,80,1)
j=1
for (i in 1:length(avged_main10g)){
  avg <- mean(zeroed_main10g[j:(j+4)])
  avged_main10g[i] <- avg
  j=j+5
}
timetopeak_period <- zeroed_main10g[110:180]
max_main <- which.max(timetopeak_period)
peak <- timetopeak_period[max_main]
time10 <- (10+max_main)*0.1

timetopeak_period <- avged_main10g[24:36]
max_main <- which.max(timetopeak_period)
peak10 <- timetopeak_period[max_main]
#time10 <- (4+max_main)*0.5

half_max <- peak10/2
half_max_period <- avged_main10g[32:80]
half_timepoint <- which.min(abs(half_max_period - half_max))
half <- half_max_period[half_timepoint]
half_time <- (12+half_timepoint)*0.5
timetohalf10 <- half_time-time10

filename_main <- "prep1_nerve1_muscle2_20Hz_green.txt"
intensity_main <- read.table(filename_main, header=TRUE)
fmain20_main <- intensity_main$main
f_main20 <- mean(intensity_main$main[50:80])
f_main20
deltaf_main <- (intensity_main$main-f_main20)/f_main20
deltaf_main20 <- (intensity_main$main-f_main20)/f_main20
min_main <- which.min(deltaf_main)
zeroed_main20g <- deltaf_main
for (i in 1:length(deltaf_main)){
  zeroed_main20g[i] <- deltaf_main[i]-deltaf_main[min_main]
}
avged_main20g <- seq(1,80,1)
j=1
for (i in 1:length(avged_main20g)){
  avg <- mean(zeroed_main20g[j:(j+4)])
  avged_main20g[i] <- avg
  j=j+5
}

timetopeak_period <- zeroed_main20g[110:180]
max_main <- which.max(timetopeak_period)
peak <- timetopeak_period[max_main]
time20 <- (10+max_main)*0.1

timetopeak_period <- avged_main20g[24:36]
max_main <- which.max(timetopeak_period)
peak20 <- timetopeak_period[max_main]
#time20 <- (4+max_main)*0.5

half_max <- peak20/2
half_max_period <- avged_main20g[34:80]
half_timepoint <- which.min(abs(half_max_period - half_max))
half <- half_max_period[half_timepoint]
half_time <- (12+half_timepoint)*0.5
timetohalf20 <- half_time-time20

filename_main <- "prep1_nerve1_muscle2_40Hz_green.txt"
intensity_main <- read.table(filename_main, header=TRUE)
fmain40_main <- intensity_main$main
f_main40 <- mean(intensity_main$main[50:80])
f_main40
deltaf_main <- (intensity_main$main-f_main40)/f_main40
deltaf_main40 <- (intensity_main$main-f_main40)/f_main40
min_main <- which.min(deltaf_main)
zeroed_main40g <- deltaf_main
for (i in 1:length(deltaf_main)){
  zeroed_main40g[i] <- deltaf_main[i]-deltaf_main[min_main]
}
avged_main40g <- seq(1,80,1)
j=1
for (i in 1:length(avged_main40g)){
  avg <- mean(zeroed_main40g[j:(j+4)])
  avged_main40g[i] <- avg
  j=j+5
}
timetopeak_period <- zeroed_main40g[110:180]
max_main <- which.max(timetopeak_period)
peak <- timetopeak_period[max_main]
time40 <- (10+max_main)*0.1

timetopeak_period <- avged_main40g[24:36]
max_main <- which.max(timetopeak_period)
peak40 <- timetopeak_period[max_main]
#time40 <- (4+max_main)*0.5

half_max <- peak40/2
half_max_period <- avged_main40g[34:80]
half_timepoint <- which.min(abs(half_max_period - half_max))
half <- half_max_period[half_timepoint]
half_time <- (12+half_timepoint)*0.5
timetohalf40 <- half_time-time40

zeroed_main_avg <- array(c(zeroed_main1g, zeroed_main2g, zeroed_main5g, zeroed_main10g, zeroed_main20g, zeroed_main40g, zeroed_main60g),dim = c(1000,4))
m0 <- c("zeroed_main5", "zeroed_main10", "zeroed_main20", "zeroed_main40")
colnames(zeroed_main_avg) <- m0
write.table(zeroed_main_avg, "190819_prep1_nerve2_muscle1_r13_green.txt")
f_main_list <- c(f_main5,f_main10,f_main20, f_main40)
write.table(f_main_list, "190819_prep1_nerve2_muscle1_r13_green_bg.txt")
time_list <- c(time5,time10,time20, time40)
time_list
write.table(time_list, "190819_prep1_nerve2_muscle1_r13_green_time.txt")
peak_list <- c(peak5,peak10,peak20, peak40)
peak_list
write.table(peak_list, "190819_prep1_nerve2_muscle1_r13_green_peak.txt")
timetohalf_list <- c(timetohalf5,timetohalf10,timetohalf20, timetohalf40)
timetohalf_list
write.table(timetohalf_list, "190819_prep1_nerve2_muscle1_r13_green_timetohalf.txt")

```


Figure 2 3 4 combined analysis
```{r jRCaMP1b muscle1}

#note that this script can be used for all combined analysis in Fig2,3 and 4 by changing the files names to reflect different data sets.

#create matrices to store values for peak df/f (f_max), background fluoresence (f_bg), time to peak (timep), and recovery time to half maximal (timeh) for jRCaMP1b at muscle 1 for ReePA+(wt)
f_max = matrix( c(2, 4), nrow = 24, ncol = 7)
f_bg = matrix( c(2, 4), nrow = 24, ncol = 7)
timep = matrix( c(2, 4), nrow = 24, ncol = 7)
timeh = matrix( c(2, 4), nrow = 24, ncol = 7)

#read in txt files created in individual scripts that contain these values at each frequency. and populate the matrices created above with these values. N=3 shown here.
filename_main <- "190806_prep1_nerve1_muscle1_red_peak.txt"
intensity_main <- read.table(filename_main, header=TRUE)
filename_bg <- "190806_prep1_nerve1_muscle1_red_bg.txt"
intensity_bg <- read.table(filename_bg, header=TRUE)
filename_timep <- "190806_prep1_nerve1_muscle1_red_time.txt"
intensity_timep <- read.table(filename_timep, header=TRUE)
filename_timeh <- "190806_prep1_nerve1_muscle1_red_timetohalf.txt"
intensity_timeh <- read.table(filename_timeh, header=TRUE)
for (i in 1:7){
  f_max[1,i] <- intensity_main[i,1]
  f_bg[1,i] <- intensity_bg[i,1]
  timep[1,i] <- intensity_timep[i,1]
  timeh[1,i] <- intensity_timeh[i,1]
}

filename_main <- "190807_prep1_nerve1_muscle1_red_peak.txt"
intensity_main <- read.table(filename_main, header=TRUE)
filename_bg <- "190807_prep1_nerve1_muscle1_red_bg.txt"
intensity_bg <- read.table(filename_bg, header=TRUE)
filename_timep <- "190807_prep1_nerve1_muscle1_red_time.txt"
intensity_timep <- read.table(filename_timep, header=TRUE)
filename_timeh <- "190807_prep1_nerve1_muscle1_red_timetohalf.txt"
intensity_timeh <- read.table(filename_timeh, header=TRUE)
for (i in 1:7){
  f_max[2,i] <- intensity_main[i,1]
  f_bg[2,i] <- intensity_bg[i,1]
  timep[2,i] <- intensity_timep[i,1]
  timeh[2,i] <- intensity_timeh[i,1]
}

filename_main <- "190808_prep1_nerve1_muscle1_red_peak.txt"
intensity_main <- read.table(filename_main, header=TRUE)
filename_bg <- "190808_prep1_nerve1_muscle1_red_bg.txt"
intensity_bg <- read.table(filename_bg, header=TRUE)
filename_timep <- "190808_prep1_nerve1_muscle1_red_time.txt"
intensity_timep <- read.table(filename_timep, header=TRUE)
filename_timeh <- "190808_prep1_nerve1_muscle1_red_timetohalf.txt"
intensity_timeh <- read.table(filename_timeh, header=TRUE)
for (i in 1:7){
  f_max[3,i] <- intensity_main[i,1]
  f_bg[3,i] <- intensity_bg[i,1]
  timep[3,i] <- intensity_timep[i,1]
  timeh[3,i] <- intensity_timeh[i,1]
}


#print matrices to make sure all values look correct. 2s and 4s will be in unused rows, and columns 5,6 and 7 are not used in this particular script.
f_max
f_bg
timep
timeh

#write the matrices to csv files so they can be opened in excel and values put into Prism for graphs and analyses.
write.csv(f_max, "fmax.csv")

write.csv(f_bg, "fbg.csv")

write.csv(timep, "fmax_timep.csv")

write.csv(timeh, "fmax_timeh.csv")

```


```{r GCaMP210ER muscle 1}

#create matrices to store values for peak df/f (f_max), background fluoresence (f_bg), time to peak (timep), and recovery time to half maximal (timeh) for ER-GCaMP-210 at muscle 1 for ReePA+(wt)
f_maxg = matrix( c(2, 4), nrow = 24, ncol = 7)
f_bgg = matrix( c(2, 4), nrow = 24, ncol = 7)
timepg = matrix( c(2, 4), nrow = 24, ncol = 7)
timehg = matrix( c(2, 4), nrow = 24, ncol = 7)

#read in txt files created in individual scripts that contain these values at each frequency. and populate the matrices created above with these values. N=3 shown here.
filename_main <- "190806_prep1_nerve1_muscle1_green_peak.txt"
intensity_main <- read.table(filename_main, header=TRUE)
filename_bg <- "190806_prep1_nerve1_muscle1_green_bg.txt"
intensity_bg <- read.table(filename_bg, header=TRUE)
filename_timep <- "190806_prep1_nerve1_muscle1_green_time.txt"
intensity_timep <- read.table(filename_timep, header=TRUE)
filename_timeh <- "190806_prep1_nerve1_muscle1_green_timetohalf.txt"
intensity_timeh <- read.table(filename_timeh, header=TRUE)
for (i in 1:7){
  f_maxg[1,i] <- intensity_main[i,1]
  f_bgg[1,i] <- intensity_bg[i,1]
  timepg[1,i] <- intensity_timep[i,1]
  timehg[1,i] <- intensity_timeh[i,1]
}

filename_main <- "190807_prep1_nerve1_muscle1_green_peak.txt"
intensity_main <- read.table(filename_main, header=TRUE)
filename_bg <- "190807_prep1_nerve1_muscle1_green_bg.txt"
intensity_bg <- read.table(filename_bg, header=TRUE)
filename_timep <- "190807_prep1_nerve1_muscle1_green_time.txt"
intensity_timep <- read.table(filename_timep, header=TRUE)
filename_timeh <- "190807_prep1_nerve1_muscle1_green_timetohalf.txt"
intensity_timeh <- read.table(filename_timeh, header=TRUE)
for (i in 1:7){
  f_maxg[2,i] <- intensity_main[i,1]
  f_bgg[2,i] <- intensity_bg[i,1]
  timepg[2,i] <- intensity_timep[i,1]
  timehg[2,i] <- intensity_timeh[i,1]
}

filename_main <- "190808_prep1_nerve1_muscle1_green_peak.txt"
intensity_main <- read.table(filename_main, header=TRUE)
filename_bg <- "190808_prep1_nerve1_muscle1_green_bg.txt"
intensity_bg <- read.table(filename_bg, header=TRUE)
filename_timep <- "190808_prep1_nerve1_muscle1_green_time.txt"
intensity_timep <- read.table(filename_timep, header=TRUE)
filename_timeh <- "190808_prep1_nerve1_muscle1_green_timetohalf.txt"
intensity_timeh <- read.table(filename_timeh, header=TRUE)
for (i in 1:7){
  f_maxg[3,i] <- intensity_main[i,1]
  f_bgg[3,i] <- intensity_bg[i,1]
  timepg[3,i] <- intensity_timep[i,1]
  timehg[3,i] <- intensity_timeh[i,1]
}


#print matrices to make sure all values look correct. 2s and 4s will be in unused rows, and columns 5,6 and 7 are not used in this particular script.
f_maxg
f_bgg
timepg
timehg

#write the matrices to csv files so they can be opened in excel and values put into Prism for graphs and analyses.
write.csv(f_maxg, "fmaxg.csv")

write.csv(f_bgg, "fbgg.csv")

write.csv(timepg, "fmaxg_timepg.csv")

```
